# A bivariate zero-inflated negative binomial model and its applications to biomedical settings

**DOI:** 10.1101/2020.03.06.977728

**Authors:** Hunyong Cho, Chuwen Liu, John S. Preisser, Di Wu

## Abstract

The zero-inflated negative binomial (ZINB) distribution has been widely used for count data analyses in various biomedical settings due to its capacity of modeling excess zeros and overdispersion. When there are correlated count variables, a bivariate model is essential for understanding their full distributional features. For this purpose, we develop a Bivariate Zero-Inflated Negative Binomial (BZINB) model that has a simple latent variable framework and parameters with intuitive interpretations. Using this model, we examine two biomedical data examples where the counts are zero-inflated—single cell RNA sequencing (scRNA-seq) data and dental caries count indices. In scRNA-seq data example, a correlation between a pair of genes is estimated after adjusting for the effects of dropout events represented by excess zeros. In the dental caries data, we analyze how the treatment with Xylitol mints affects the marginal mean and other patterns of response manifested in the two dental caries traits. An `R` package ‘`bzinb`’ is available on CRAN.

## 1. Introduction

### 1.1 Motivation

In biomedical research, count data often include a large number of zeros. For example, quite often people do not have dental caries (Preisser *and others*, 2016), and the majority of a population does not make a hospital visit during a given year (Gurmu, 1997). In omics data, either because of technological reasons related to sequencing or due to some biologoical reasons, counts are often very sparse (Risso *and others*, 2018; Van den Berge *and others*, 2018; Aldirawi *and others*, 2019). Such count data with excess number of zeros are frequently modeled using the zero-inflated Poisson (ZIP) or the zero-inflated negative binomial (ZINB) distributions (Lambert, 1992; Greene, 1994). The negative binomial distribution, of which the Poisson distribution is a limiting case, has a capacity of modeling overdispersion that accounts for heterogeneity of the incidence processes and thus is widely used in practice. Zero-inflation refers to the phenomenon where the proportion of zeros are greater than is expected by the corresponding reference distribution.

However, these univariate models only provide insights about marginal distributions and do not inform the joint characterization of two dependent count variables. For example, when the association of two genes is of interest, a bivariate model is needed to effectively estimate the dependence. On the other hand, in caries and other clinical trials, a joint model may be beneficial when univariate analyses are less efficient for comparing marginal mean outcomes between treatment groups. As in the choice of MANOVA versus multiple ANOVAs for the hypothesis testing of continuous random variables, multivariate count models can control type-I error in a more efficient way than multiple testing based on univariate ZIP or ZINB models. To meet these different purposes, we develop a flexible bivariate zero-inflated negative binomial model that characterizes the joint distribution of two correlated count random variables.

The two motivating examples are used to illustrate the utility of the proposed bivariate model for count data in biomedical research. The first one involves the measurement of dependence of two genes in single cell RNA sequencing (scRNA-seq) data that have a significant amount of dropouts, which are represented by excess zeros under model assumptions. The pairwise dependencies are important as they serve as building blocks for identifying pathways. However, possible dropout events obfuscate measuring dependence of gene pairs in the absence of dropout. For this task, we measure dependence after controlling for the dropout events through a bivariate zero-inflation model. The second example is characterizing the distribution of the number of dental caries detected on two different tooth surfaces. A treatment—Xylitol mints—may not only affect the mean count for both traits to varying degrees but also may modify the structure of the joint distribution. A bivariate model is needed to adequately characterize the joint distribution and estimate marginal effects on outcomes with possibly improved precision. More details of the two examples are given in the following subsections.

### 1.2 Measuring genewise dependence in scRNA-seq data

Single cell RNA sequencing (scRNA-seq) is a high throughput sequencing technology that profiles gene expression at a cell’s resolution (Kolodziejczyk *and others*, 2015). This is in contrast to bulk RNA sequencing (RNA-seq), where a group of cells are sequenced altogether and consequently no cell-level information is available in data. As a price for cell-level resolution, scRNA-seq loses some information by the so-called “dropout” phenomenon; during the sequencing steps (and the capturing steps, e.g., in 10X sequencing platform) of scRNA-seq, a large amount of RNAs are undetected. Consequently, the observed count data include a greater number of zeros than would be expected given the number of molecules sequenced and our *a priori* knowledge of transcription rates at individual loci (Risso *and others*, 2018; Hicks *and others*, 2017; Huang *and others*, 2018). In contrast, in a bulk RNA-seq, excess zeros are less frequently observed (Hicks *and others*, 2017). For these reasons, negative binomial models have been extensively used for bulk RNA-seq data (Love *and others*, 2014; Robinson *and others*, 2010), and ZINB models are typically used for scRNA-seq data (Van den Berge *and others*, 2018; Risso *and others*, 2018). Although recent technologies such as the unique moecular identifiers (UMI) effectively remove the dropout events, the problem still persists in data obtained from many of the non-UMI-based platforms.

Statistical inferences at both individual gene level (Iacono *and others*, 2019; Yu, 2018) and gene set level, e.g., pathways, can be misleading without considering the excess zeros caused by dropouts. Inference of gene-gene dependence, e.g., the correlation-based method, has been widely used in pathway analysis of bulk RNA-seq data (Zhang and Horvath, 2005) and in recent scRNAseq data analyses (Iacono *and others*, 2019; Yu, 2018; Pont *and others*, 2019; Van Dijk *and others*, 2018; Eraslan *and others*, 2019). However, the conventional Pearson correlation of two genes with significant dropouts in the scRNAseq may not properly reflect the underlying gene-gene dependence.

For example, a pair of genes, of which expressions are highly correlated in the absence of dropouts, would have an attenuated correlation, based on the observed data, if only one of the genes have a large amount of dropouts. On the other hand, a pair of uncorrelated genes would have higher correlation, when both genes have dropouts in a substantial portion of the sample. The systematic bias will not vanish without adjusting for the effects of the dropout events, regardless of the dependence measure such as Pearson correlation and mutual information.

Two strategies have been considered to address the bias in scRNA-seq data. Imputation methods (Li and Li (2018); Eraslan *and others* (2019); Peng *and others* (2019)) aim to provide expression levels free of the excess zeros by imputing them. While imputation methods are versatile in that they provide ready-to-use data, they are not deterministic, having different results for every implementation. The other strategy is estimation of the count distribution (Huang *and others*, 2018; Wang *and others*, 2018). Once having obtained information about the distribution of the expressions before dropouts, one can do downstream analyses such as measuring the dependence of the before-dropout expressions. However, many of the methods taking this approach focus on modeling marginal distributions and they do not explicitly posit dependence structure between two genes.

Our proposed method, to be introduced in Section 1.4, takes the distribution estimation approach where a bivariate distribution explicitly addresses the dependence structure. Specifically, our method is built on a bivariate generalization of the zero-inflated negative binomial (ZINB) model, where the dropout probabilities are modeled using the zero-inflation parameters. In this approach the full joint distribution is estimated using a bivariate count model with zero-inflation and the underlying distribution before dropouts is uncovered to measure the dependence.

### 1.3 Characterizing the joint count distribution of two dental caries traits

The second example is about describing the pattern of the number of dental caries occurring at two different tooth surface types in a randomized clinical trial (the Xylitol for Adult Caries Trial, or X-ACT) (Bader *and others*, 2013). In one of its secondary studies, the effect of Xylitol mints on the incidence of three different dental caries outcomes were examined with univariate models according to the type of tooth surface: smooth-surface caries, occlusal-surface caries, and proximal-surface caries (Ritter *and others*, 2013). There are often a great amount zero counts for dental caries and the ZINB model is thus frequently used in the literature (Preisser *and others*, 2017), as exemplified by this study. Of note, the excess zeros in this example have a different nature than those in scRNA-seq data in the sense that those zeros more likely represent lack of incidence from the “non-susceptible” population and are not induced by some unwanted events such as dropouts in the scRNA-seq setting.

However, the original analysis can only tell us about the marginal distribution of each trait and does not provide any information of their joint structure. For example, Xylitol mints may have lowered the correlation between the proximal- and smooth-surface caries by reducing the concordant pairs (zero for both or non-zero for both) while increasing the discordant pairs (zero for one and non-zero for the other). A joint analysis may give additional clues for identifying the mechanism of Xylitol mints and can only be obtained through joint modeling of the traits. We analyze the joint distribution of a pair of traits using our bivariate extension of the ZINB distribution.

The bivariate models could also be used to boost power in marginal analyses, since the information contained in one variable could be borrowed in making inferences on the other. For instance, in testing the group difference of mean incidence rates on smooth surfaces, the joint incidence rate of smooth and proximal surfaces could be estimated with a bivariate model and the group differences could be tested for each margin at an enhanced power.

### 1.4 Review of existing bivariate count models and the proposed method

In consideration of building bivariate count models, it is noteworthy that there have been proposed a variety of bivariate models that fit overdispersed count data: bivariate Poisson mixture models (Gurmu and Elder, 1999), (Famoye, 2010), and (Jørgensen, 1987)), bivariate generalized Poisson models (Famoye and Consul, 1995) and copula models (Cameron *and others*, 2004). These models can be further extended to flexibly accommodate excess zeros by introducing zero-inflation parameters or composing hurdle models. For a comprehensive survey of bivariate count models, refer to Cameron and Trivedi (2013) and Chou and Steenhard (2011).

Of a plethora of the proposed models in the literature, many of the bivariate Poisson mixture models and bivariate generalized Poisson models take overly complicated forms, they do not have simple marginal distributions (e.g., GBIVARNB model in Gurmu and Elder (1999)), and their parameters are hard to interpret and/or computationally expensive to estimate. Copula-based bivariate models can be alternatives to the mixture models, but they depend on the underlying copula models and can be difficult to interpret.

Many existing bivariate negative binomial models are primarily designed for modeling marginal means rather than pairwise dependence. For example, Gurmu and Elder (1999) discussed a bivariate negative binomial distribution (BIVARNB), but their model is specified by only four parameters, which may not provide sufficient flexibility to delineate diverse distributional structure. For such a bivariate joint distribution, four parameters are needed to specify the first two marginal moments of each of the two variables, while another parameter is needed solely for modeling the dependence. Subsequently Wang (2003) extended BIVARNB to a zero-inflated BIVARNB regression setting. In this model, zero-inflation is dictated by a single parameter, implying that when one variable either drops out or not, the other variable behaves exactly the same, which may not be the case for scRNA-seq data; one gene can drop out, while the other does not in a sample. Instead, it is possible to have three free parameters for the full joint zero-inflation probability structure (Li *and others*, 1999).

We propose a bivariate zero-inflated negative binomial model with eight parameters: five parameters for the negative binomial part and another three free parameters for the zero-inflation part. This model allows analyzing the dependence of two zero-inflated count variables parametrically but with more flexibility than existing models. Specifically, five parameters of our proposed model characterize all moments of the first two orders before zero-inflation, and the three zero-inflation parameters model the dropouts or the membership of non-susceptible groups with full flexibility.

Besides the flexibility and the provision of the dependence measure, the proposed BZINB model has the following features. The parameters have simple latent variable interpretations, the joint distribution can be marginalized into the corresponding univariate ZINB distributions, and the model can be easily reduced to a non-zero-inflated model, or BNB, by dropping the zero-inflation parameters.

The rest of the paper is organized as follows. In Section 2, we describe how the model is constructed based, successively, on a Bivariate Negative Binomial model and a Bivariate Zero-inflated Negative Binomial model. We present the maximum likelihood estimator using the expectation-maximization (EM) algorithm in Section 3. In Section 4, we illustrate how well the model fits data and how the model-based dependence measure behaves in contrast to naive measures using mouse paneth scRNA-seq data and study the performance of the dependence measure via simulations. In Section 5, we analyze the dental caries clinical trial data with the BZINB model to characterize the joint distribution of two dental caries traits, and illustrate the use of the model in testing the group differences in the marginal means of the two traits. In Section 6, we address limitations of the models and discuss potential extensions. Section 7 provides software information, and Section 8 lists the Supplementary Materials.

## 2. The model

### 2.1 A Bivariate Negative Binomial Model

In constructing the BZINB model, to induce dependence and zero-inflation, layers of latent variables were used as in Kocherlakota and Kocherlakota (1992) and Li *and others* (1999). We first introduce a simpler model, the Bivariate Negative Binomial (BNB) model in this subsection, and then generalize it to Bivariate Zero-Inflated Negative Binomial (BZINB) model in Subsection 2.2. One of the key assumptions about the dependence structure of BNB (and BZINB) is that the mean parameters of two Poisson random variables are gamma random variables that share a common gamma random variable. Let *R*_*j*_ ∼ *Gamma*(*α*_*j*_, *β*) for *j* = 0, 1, 2, where *α*_*j*_ and *β* are the shape and scale parameters, respectively. Then (*R*_0_ + *R*_1_, *R*_0_ + *R*_2_) is bivariate gamma distributed, denoted as *BGamma*(*α*_0_, *α*_1_, *α*_2_, *β*). To account for heterogeneous scales of the two Poisson mean variables, we introduce an additional parameter *δ* ∈ ℝ^+^. Then, a pair (*X*_1_, *X*_2_) of Poisson variables with means (*R*_0_ + *R*_1_, *δ*(*R*_0_ + *R*_2_)) follow a bivariate negative binomial distribution, denoted as

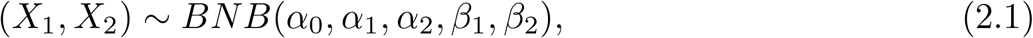

where we reparametrize (*β, δ*) as (*β*_1_, *β*_2_) = (*β, δβ*) and the observed density is given as,

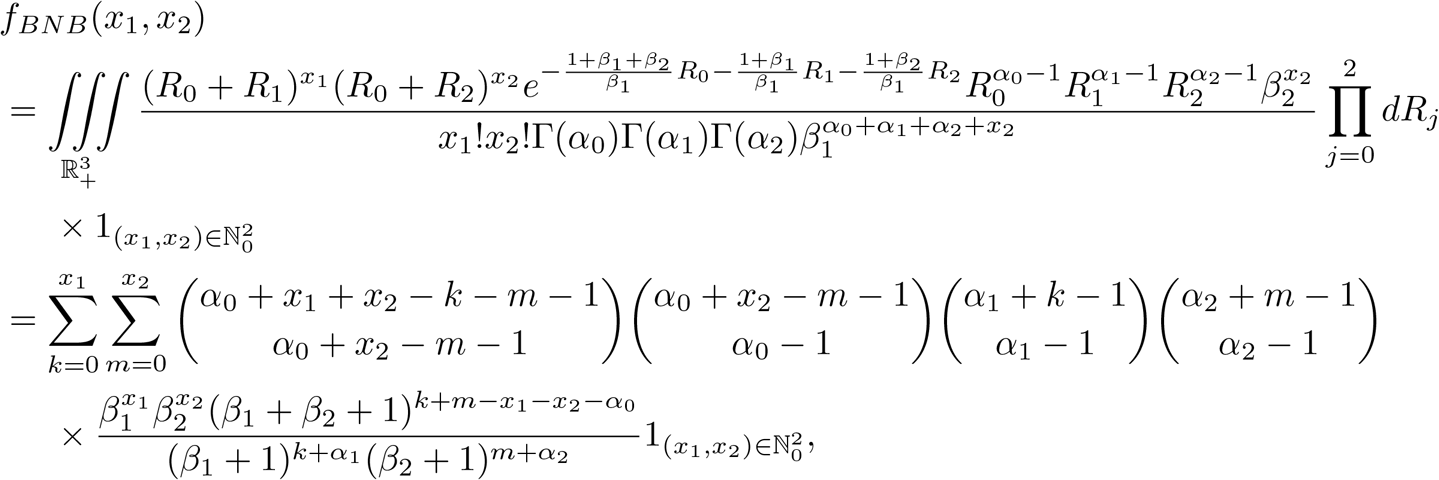

where ℝ_+_ and ℕ_0_ denote the positive real and nonnegative integer spaces, respectively, and superscripts represent the dimension of the product space. The support indicators will be omitted throughout this paper when the context is clear.

This bivariate negative binomial model (BNB) is marginally negative binomial, as we know from the construction procedure that both *X*_1_ and *X*_2_ are Poisson random variables with means marginally Gamma distributed, respectively:

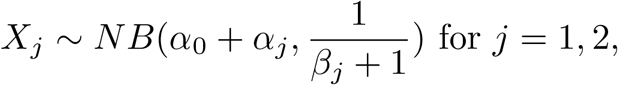

where the random variable *X* ∼ *NB*(*ν, ϕ*) can be interpreted as the minimum number of failures to have *ν* successes with probability of *ϕ* for each trial; i.e., its density is expressed as 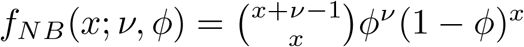.

Interpretation of the BNB parameters is straightforward: *α*_0_, *α*_1_, and *α*_2_ are the shape parameters of latent variables, where the larger *α*_0_ implies a larger amount of shared components in *X*_1_ and *X*_2_ and thus larger correlation; *β*_1_ and *β*_2_ controls the scale of *X*_1_ and *X*_2_, respectively.

Note in scRNA-seq data context, *X*_1_ and *X*_2_ may represent the *before-dropout* expression level of each of two genes in a cell in the absence of dropout events, which we rarely observe in practice.

The first two moments and the correlation of a BNB random pair are given as,

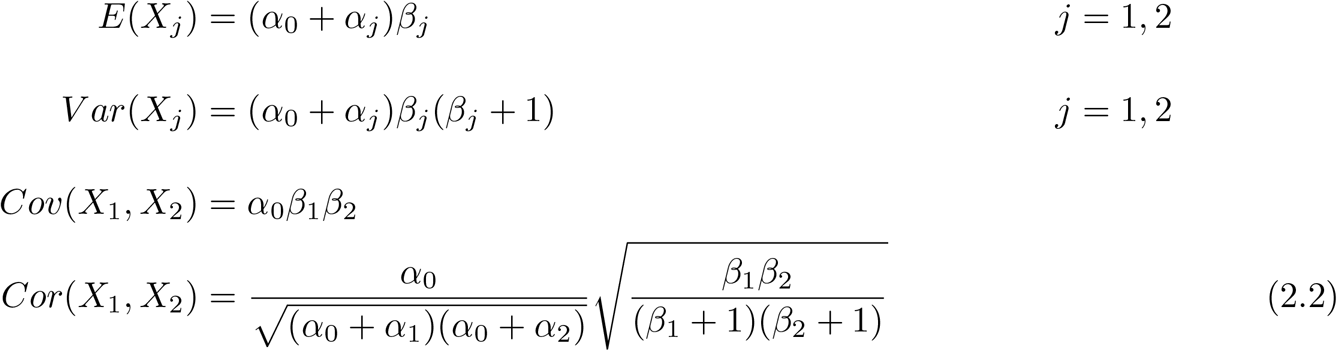

Note that this distribution only allows positive correlation. See Section 6 for more discussion.

Maher (1990) developed another bivariate negative binomial distribution that is a constrained case of BNB in a sense that the marginal means and variances are the same for both variables.

One can further generalize this BNB model into a *m*-variate negative binomial model by adding common latent gamma parameter(s) to the *m* gamma variables.

### 2.2 A Bivariate Zero-inflated Negative Binomial Model

In this subsection, we generalize BNB model to BZINB model by including zero-inflation components. Since BZINB is also a generalization of univariate zero-inflated negative binomial model (ZINB), we illustrate the construction of univariate ZINB model first and move to the bivariate version.

A univariate negative binomial model, *NB*(*ν, ϕ*), can be generalized to allow zero-inflation by having an additional parameter, *π*: *ZINB*(*ν, ϕ, π*). The zero-inflated negative binomial (ZINB) model has a latent variable interpretation. Let *X* follow *NB*(*ν, ϕ*) and *E* denote the zero-inflation indicator having 1 with probability of *π* and 0 otherwise, independently of *X*. Then *Y* ≡ (1−*E*)*X* follows *ZINB*(*ν, ϕ, π*) with the density of *f*_*ZINB*_(*y*; *ν, ϕ, π*) = (1 − *π*)*f*_*NB*_(*y*; *ν, ϕ*) + *πζ*(*y*), where *ζ*(*a*) ≡ 1_(*a*=0)_.

Similarly, a multivariate zero-inflated random variable can be constructed using a latent variable that follows the multivariate Bernoulli distribution as in the Poisson case (Li *and others*, 1999). For a bivariate distribution, suppose we have a random vector ***E*** ≡ (*E*_1_, *E*_2_, *E*_3_, *E*_4_)^⊤^ ∼ *MN* (1, ***π***), where *MN* (1, ***π***) denotes the multinomial distribution with a single trial and an associated probability of ***π*** ≡ (*π*_1_, *π*_2_, *π*_3_, *π*_4_)^⊤^ with **1**^⊤^***π*** = 1. Now the bivariate zero-inflated negative binomial distribution (BZINB) can be formulated as:

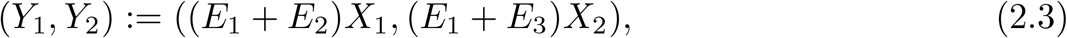

where (*X*_1_, *X*_2_) ∼ *BNB*(*α*_0_, *α*_1_, *α*_2_, *β*_1_, *β*_2_) and *E*_1_, *E*_2_, *E*_3_ and *E*_4_ are the indicators of observing both *X*_1_ and *X*_2_, only *X*_1_, only *X*_2_, and none of them, respectively. We say (*Y*_1_, *Y*_2_) ∼ *BZINB*(*α*_0_, *α*_1_, *α*_2_, *β*_1_, *β*_2_, *π*_1_, *π*_2_, *π*_3_). A simpler model with a restriction of *π*_2_ = *π*_3_ = 0 can also be considered as in Wang (2003).

The density of a BZINB variable is

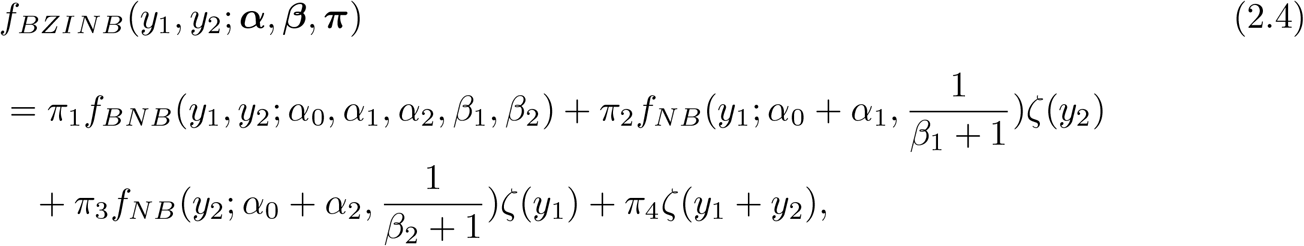

where ***α*** = (*α*_0_, *α*_1_, *α*_2_)^⊤^, ***β*** = (*β*_1_, *β*_2_)^⊤^, and ***π*** = (*π*_1_, *π*_2_, *π*_3_, *π*_4_)^⊤^ with **1**^⊤^***π*** = 1.

Here, the parameters ***α*** and ***β*** have the same interpretation as in BNB but in the presence of dropouts, and ***π*** indicates the dropout probability, where *π*_1_, *π*_2_, *π*_3_, and *π*_4_ are the probability that none, *Y*_2_ only, *Y*_1_ only, and both were dropped out, respectively.

In scRNA-seq data, *Y*_1_ and *Y*_2_ are the *observed* number of expressions for each of two genes in a cell. The term *observed* was used in contrast to *before-dropout* in a sense that an unobserved subset of the zeros are excess zeros due to dropouts.

This BZINB distribution is marginally ZINB, since the latent random variables, *X*_1_ and *X*_2_, are marginally negative binomial random variables (from Subsection 2.1) with probabilities of being observed, *π*_1_ + *π*_2_ and *π*_1_ + *π*_3_, respectively:

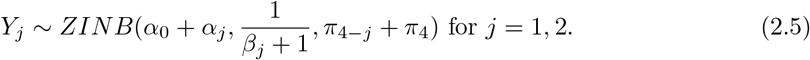

The first two moments of a BZINB pair are given as,

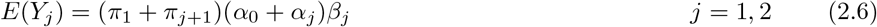

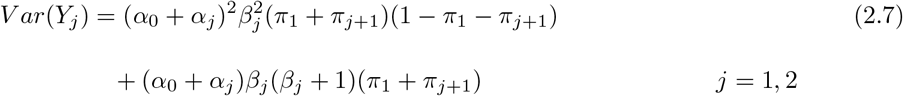

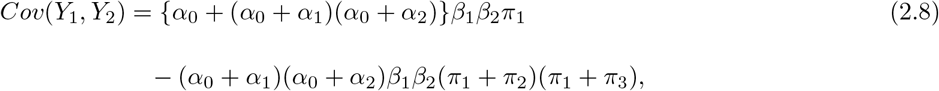

and the correlation *ρ*(*Y*_1_, *Y*_2_) is not further simplified than 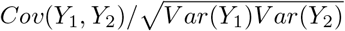.

When dropouts are unwanted and need to be adjusted for, then the underlying correlation *ρ*^*^ of *Y*_1_ and *Y*_2_ under BZINB model is simply the correlation of *X*_1_ and *X*_2_ (Equation (2.2)), which is

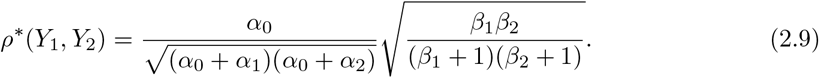

## 3. Estimation

With the natural interpretation of BZINB model as layers of latent variables, one can estimate the parameters by the expectation-maximization (EM) algorithm.

The complete density is given as,

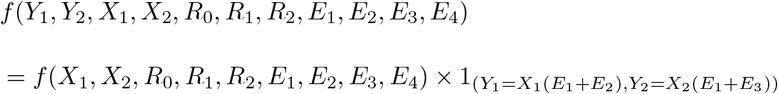

with

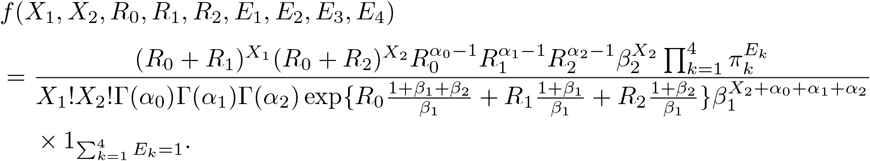

Thus, the full individual log-likelihood for the *i*th entry, or the *i*th cell, is

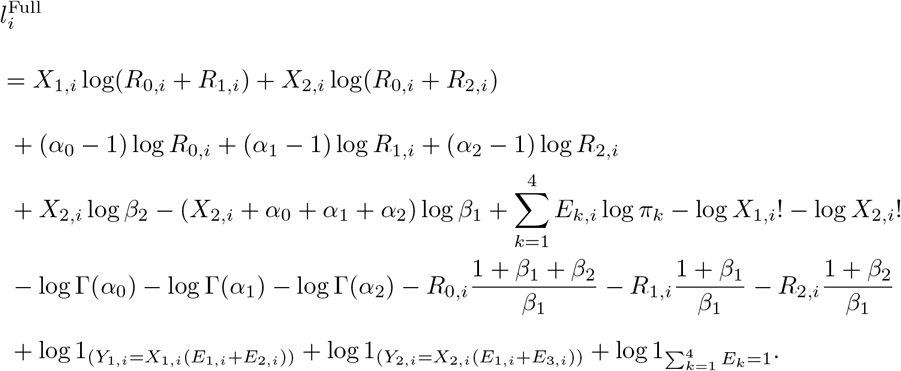

The expected full log-likelihood conditional on the observed data is linear in *E*[*R*_*j,i*_|*Y*_1,*i*_, *Y*_2,*i*_; ***θ***], *E*[log(*R*_*j,i*_|*Y*_1,*i*_, *Y*_2,*i*_; ***θ***)], *E*[*E*_*k,i*_|*Y*_1,*i*_, *Y*_2,*i*_; ***θ***], and *E*[*X*_2,*i*_|*Y*_1,*i*_, *Y*_2,*i*_; ***θ***], where ***θ*** ≡ (***α***^⊤^, ***β***^⊤^, ***π***^⊤^)^⊤^, *j* = 0, 1, 2 and *k* = 1, 2, 3, 4. The formulae of the components are given in Web Appendix A.

As the likelihood is the product of functions convex with respect to each of the parameters, the maximization can be achieved by solving a system of score equations. The individual scores are given as:

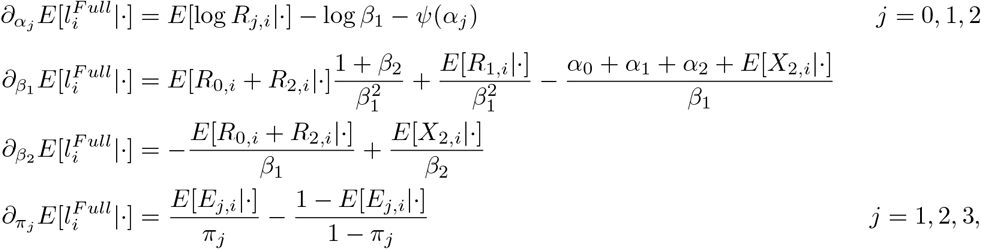

where the conditioning arguments (***Y***_1_, ***Y***_2_; ***θ***) are suppressed as (·) and can be replaced with (*Y*_1,*i*_, *Y*_2,*i*_; ***θ***) where we assume a sample of independent entries, ***Y***_*l*_ denotes (*Y*_*l*,1_, …, *Y*_*l,n*_)^⊤^ for *l* = 1, 2, *n* is the sample size, and *∂*_*a*_*b* denotes the partial derivative of *b* with respect to *a*.

At the *k* + 1st iteration of the EM algorithm, we get ***θ***^(*k*+1)^ by solving the score equations 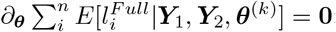:

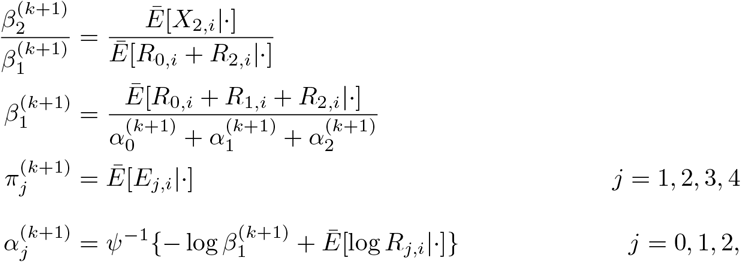

where *Ē* [*A*|·] denotes the empirical average of the conditional expectations, i.e., 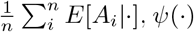 is the digamma function, and the conditioning arguments (***Y***_1_, ***Y***_2_, ***θ***^(*k*)^) are again suppressed. The equations can be solved by solving the following through Newton-Raphson algorithm:

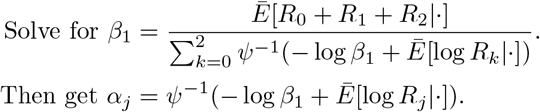

After iterations enough to observe convergence, the final updated parameter values serve as the maximum likelihood estimate.

The standard error of the maximum likelihood parameter estimates can be calculated using observed information. In Web Appendix B, detailed formulae are given, and simulations illustrating the accuracy of standard error estimation are included in Section 4.3.

## 4. Measuring gene-gene correlations accounting for dropout events by the BZINB model

### 4.1 Model comparison using the mouse paneth data

In this section, we show how the BZINB model fits a scRNA-seq data set compared to its nested models (in Subsection 4.1), present how model-based dependence measures can be different from naive measures (in Subsection 4.2), and study the asymptotic behavior of the estimator through simulations in Subsection 4.3. The data were collected from paneth cells of a C57Bl6 mouse with a Sox9 gene knockout. The Fluidigm C1 system was used to capture single cells and generate Illumina libraries using manufacutrers’ protocols. Illumina NextSeq sequencing platform was used for paired end sequencing. Reads per cell were demultiplexed using mRNASeqHT_demultiplex.pl, a script provided by Fluidigm. Low quality base calls and primers were removed using Trimmomatic (Bolger *and others*, 2014) and poly-A tails were removed using a custom perl script. Reads were aligned to the mouse genome (mm9) using STAR (https://academic.oup.com/bioinformatics/article/29/1/15/272537) and read per gene were counted using htseq-count (https://www.ncbi.nlm.nih.gov/pmc/articles/PMC4287950).

The data are composed of 23,425 genes for 800 cells, where all the cells came from a single mouse and have the same cell type. Over 90% of genes have more than 90% zero counts and the average proportion of zero counts for a gene is 97.3% in the data. We perform zero-inflation test to verify the existence of zero-inflation, since high proportion of zeros may not necessarily mean zero-inflation. The likelihood ratio test comparing the NB and ZINB distribution is used for this test with a 50:50 mixture 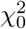 and 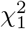 distributions as the reference distribution, where 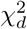 denotes a central *χ*^2^ distribution with *d* degrees of freedom (Shapiro, 1985). Figure 1 provides the histogram of p-values associated with the zero-inflation test. Without zero-inflation, the histogram should be even over the [0, 0.5] interval. However, a hike on the left is observed indicating the existence of zero-inflation, and thus overall, the use of the BZINB model is suggested over the BNB model.

**Fig. 1.**
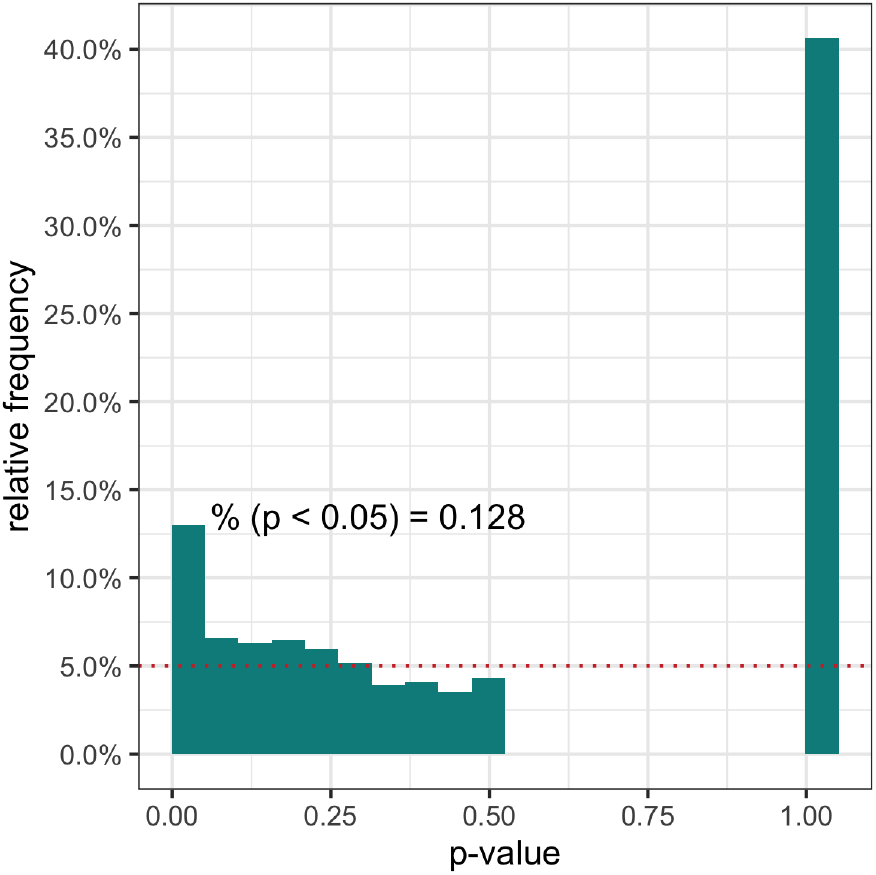
The histogram of the likelihood ratio test for zero-inflation. The reference *p*−value distribution under the null model is uniform over [0, 0.5] and a point mass 0.5 at 1, where the uniform density corresponds to 5% relative frequency line with the width of bins being 0.05.

We compare four nested models: BZINB, BNB, bivariate zero-inflated Poisson (BZIP), and bivariate Poisson (BP). BZIP has fixed mean values instead of latent gamma variables of BZINB, and BP further lacks zero-inflation components. The estimated densities of these models are compared with the empirical density for 50 gene pairs.

To systematically study the model performances, we performed stratified sampling of genes according to their proportion of zeros; strata H, M, L, and V include genes with ⩾ 90%, 80% to 90%, 60% to 80%, and *<* 60% zeros, respectively. Genes with ⩾ 98% of zeros and genes with extremely large expression (*>* 10, 000 counts for at least one cell) were screened out. After screening out those irregular genes, each group has 81.4%, 13.5%, 4.2%, and 0.9% of genes, respectively.

We randomly selected 5 pairs of genes from each possible combination of two strata (HH, MM, LL, VV, HM, HL, HV, ML, MV, and LV) without replacement. For each of the 50 pairs (5 pairs × 10 combinations), we estimated the parameters of the four nested models. Based on the parameter estimates, the distributions of the four models were compared. As it is not straightforward to compare the estimated model-based densities with the empirical density, we drew a random sample of size *n* = 800 from each estimated model and the resulting empirical densities were then compared (Figure 2 for several pairs and Web Figure 1 for all the 50 pairs). As we cannot preclude the chance of getting unlikely instances by doing Monte Carlo sampling, we added results of two more replicates in Web Figures 2 and 3. We furthermore illustrate the exact values of the estimated density in Figure 3 for a couple of pairs and in Web Figure 4 for all the pairs.

**Fig. 2.**
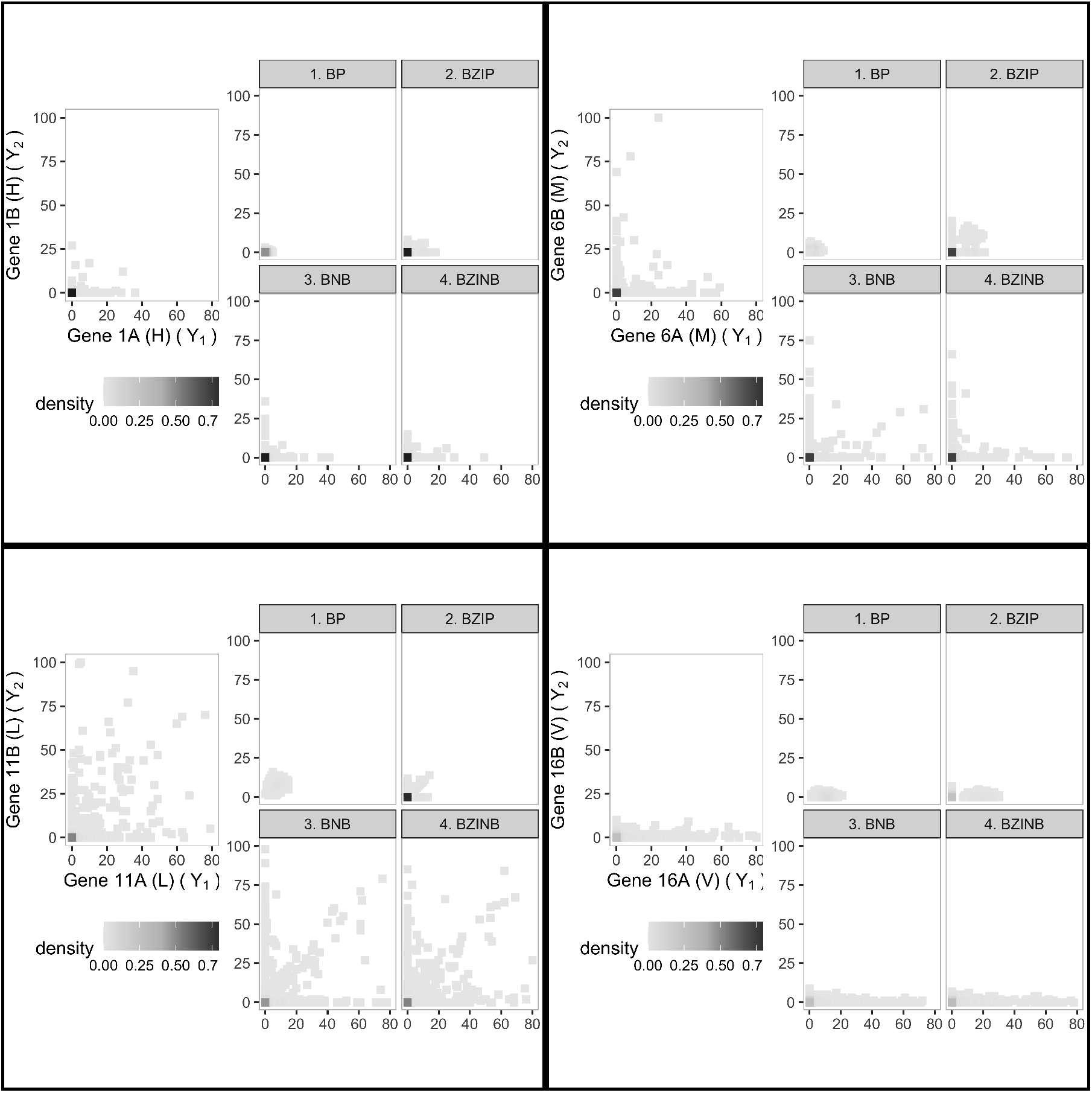
The bivariate distribution of actual and simulated mouse paneth RNA count data. Each box corresponds to the first gene pair from each stratum: HH1, MM1, LL1, VV1, where letters represent stratum with varying proportions of zeros and the numbers represent the number of the pair in each stratum. Each box has the empirical distribution (LEFT) which serves as truth for the simulations, and the four model-based simulated empirical distributions (RIGHT).

**Fig. 3.**
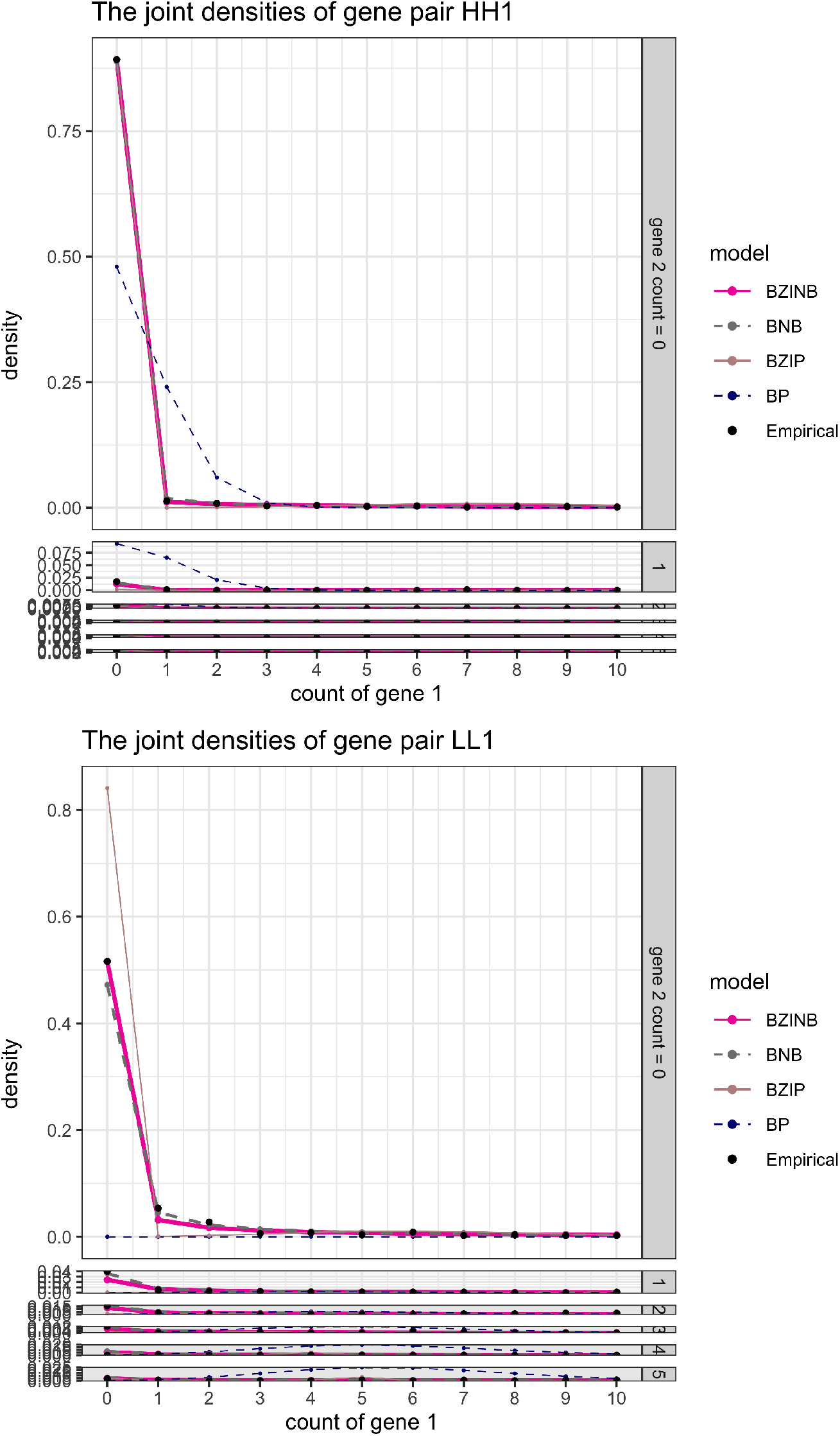
The model estimates of bivariate densities (lines) and the empirical densities (dots) of two gene pairs. The X-axis and vertical layers represent the count of the first and the second genes, respectively.

**Fig. 4.**
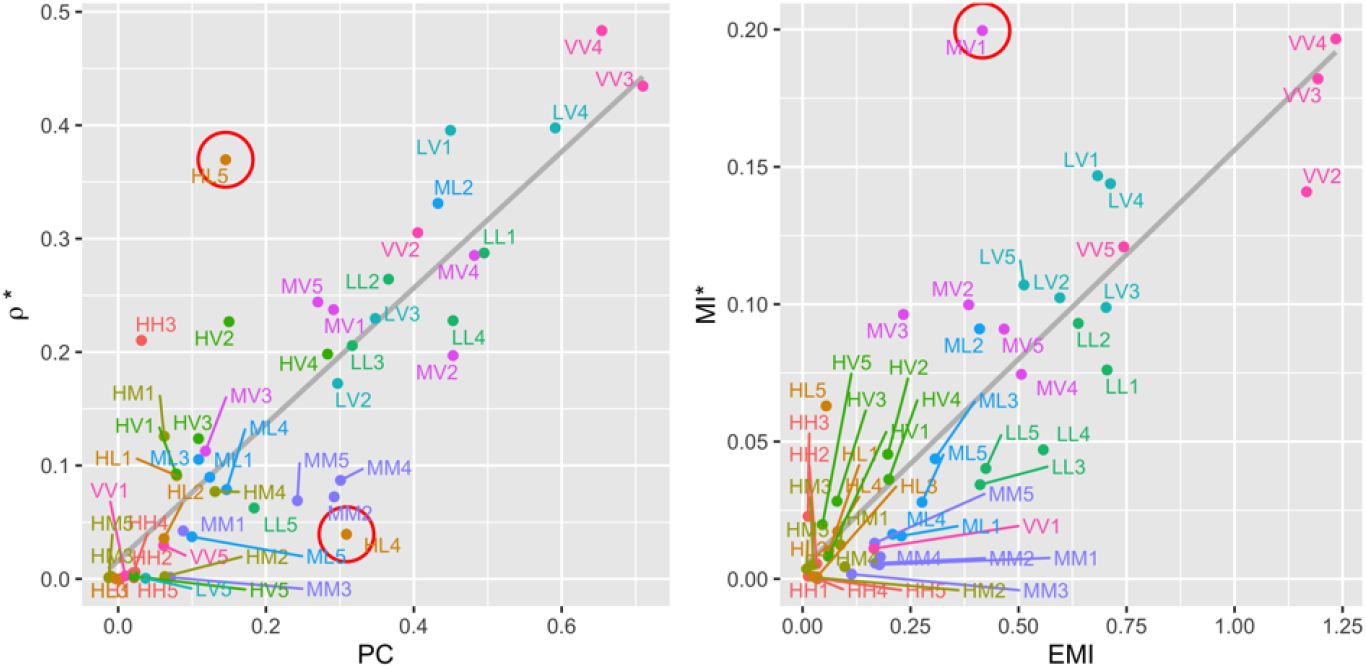
Estimated dependence measures of 50 pairs. Pearson correlation (PC) and underlying correlation estimates (*ρ*^*\**^) (LEFT). Empirical (EMI) and underlying (MI^*\**^) mutual information estimates (RIGHT).

We compare the actual and the model-based empirical distributions for the first pair from each stratum (Figure 2). The results including all 50 pairs from a single stratum or from two different strata and their replicates can be found in Web Figures 1 to 3. For any pair, the BP model obviously fails to address the overdispersion and zero-inflation, while the BZIP model could not properly mimic the overdispersion. BNB and BZINB seem to fairly mimic the true distribution in most of the pairs. The poor performances of Poisson-based models and decently good performances of BNB and BZINB models can also be seen on Figure 3.

However, when genes have some large-valued counts and many zeros at the same time either marginally or jointly, BZINB has an apparent advantage over BNB model. Often, in BNB model, nonzero count pairs are highly concentrated on the diagonal line, while nonzero counts in BZINB model are more dispersed away from the diagonal line (LL1 in Figure 2 and more examples in Web Figures 1 to 3). This can be explained by the lack of flexibility of BNB model. When data are highly zero-inflated but overdispersed at the same time, BNB is forced to have small shape parameters (*α*_*j*_, *j* = 0, 1, 2) and large scale parameters (*β*_*j*_, *j* = 1, 2) while keeping the mean of the latent Gamma variables, *E*[*R*_*j*_] = *α*_*j*_*β*_1_, close to zero. These latent Gamma variables, serving as mean parameters of Poisson variables, take on very small values most of the times and very large values with small chance. It is unlikely that both *R*_1_ and *R*_2_ have large numbers at the same time (CASE 1), but it is more frequent that *R*_0_ alone has a large number (CASE 2). Thus, the latent Poisson variables, *X*_1_ and *X*_2_, are more likely to have similarly large numbers (resulting from CASE 2) than to have significantly different nonzero numbers (resulting from CASE 1).

### 4.2 Comparison of dependence measures in the mouse paneth scRNA-seq data

When the excess zeros are believed to come from dropouts, BZINB model may uncover the underlying dependence using measures such as *ρ*^∗^ and *MI*^∗^. *MI*^∗^ is the underlying mutual information defined similarly to *ρ*^∗^ and can be estimated by first estimating the BZINB model parameters and by measuring the mutual information, 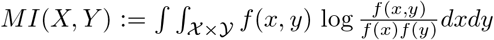, of the estimated distribution after replacing 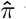 with (1, 0, 0, 0)^⊤^.

For the same 50 pairs in the previous subsection, we estimated the dependence using naive measures – Pearson correlation (PC) and empirical mutual information (EMI) – and zero-inflation adjusted measures – underlying correlation (*ρ*^∗^) and underlying MI (*MI*^∗^) based on BZINB model. Figure 4 summarizes the estimates for all the pairs. The plots of empirical distribution with estimated dependence measures for each pair are also available on Web Figure 5.

**Fig. 5.**
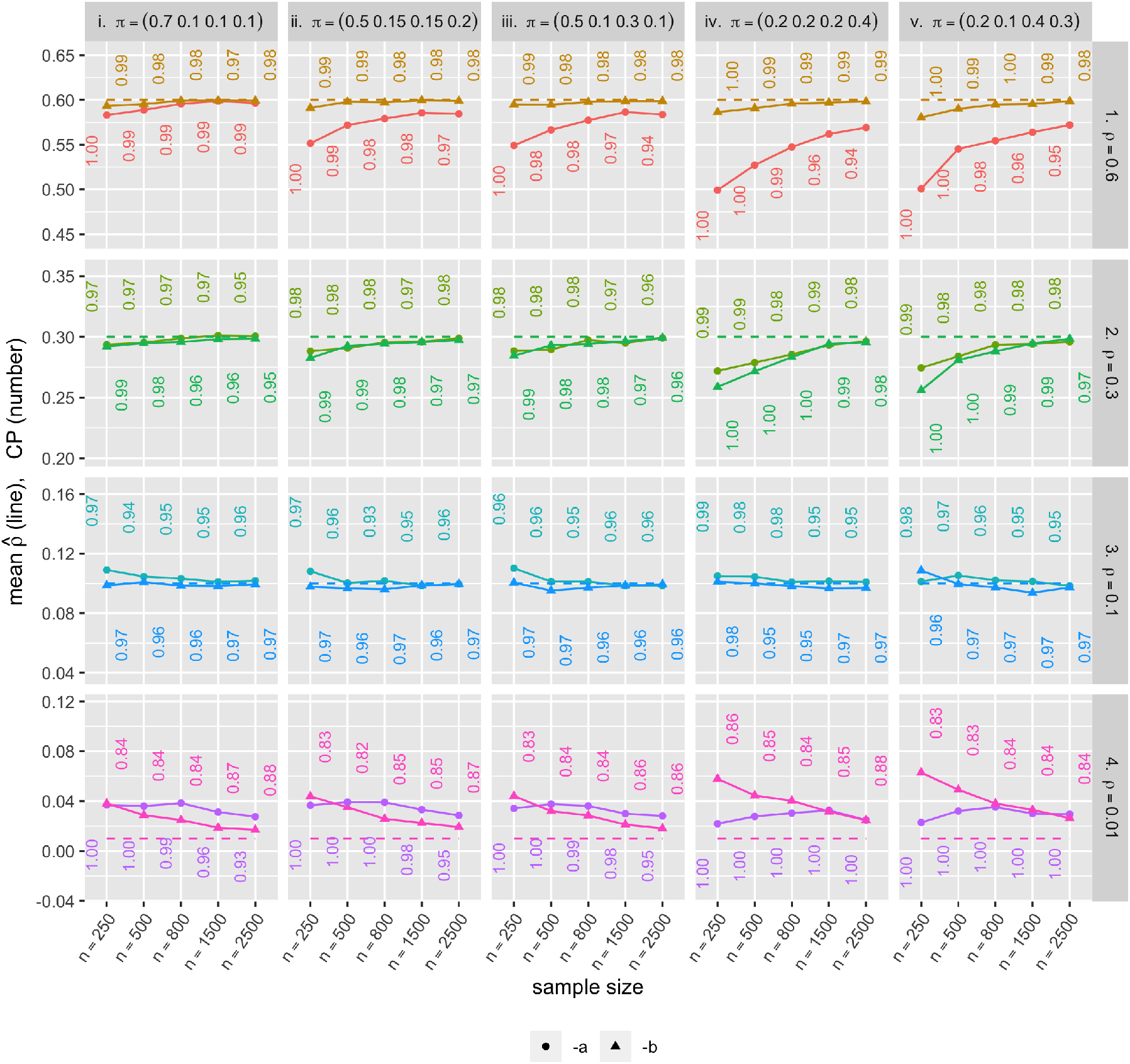
Mean parameter estimates 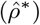 and CP (each color represents distinct simulation scenarios.)

In Figrue 4 LEFT, we see that PC and *ρ*^∗^ mostly behave in the same direction, but also that they can have values in the opposite directions (e.g., HL5 and HL4). If we judge whether two genes are correlated based on (naive) Pearson correlation (PC) with a certain threshold, say PC *>* 0.2, many genes might be missed (e.g., HL5) or falsely included (e.g., HL4).

Similar analyses can be done for MI-based measures. Both EMI and *MI*^∗^ estimates are correlated, however, there are pairs that are located away from the tendency. For example the pair MV1 has highest *MI*^∗^, while its EMI is not one of the highest. Also note that the values of *MI*^∗^ are in general less than those of EMI for scRNA-seq data. Heavy proportion of zero-zero pairs boosts naive EMI, while *MI*^∗^ removes the effects of the co-zero-inflation. These results suggest that measures that fail to identify the excess zeros caused by the dropout events may be highly misleading.

### 4.3 Evaluation of the dependence estimator based on simulation

We ran simulations to study the performance of estimators of underlying correlation and the associated standard error under finite sample size. We considered 40 distinct sets of BZINB parameter values (Table 1). Note that for each of *ρ*^*^s there are two distinct sets of parameters (***α, β***), the first (a) of which have lower ***α*** values and the second (b) of which have higher ***α*** values. For each parameter set (***α, β, π***) and for *n* = 250, 500, 800, 1500, 2500, we generated random BZINB samples of size *n, n*_sim_ = 1, 000 times.

**Table 1.** The set of parameters for simulation. Combination of (*α*_0_, *α*_1_, *α*_2_, *β*_1_, *β*_2_) and (*π*_1_, *π*_2_, *π*_3_, *π*_4_) below makes 40(= 8 × 5) sets in total.

| underlying correlation | | # | $(\alpha_0, \alpha_1, \alpha_2, \beta_1, \beta_2)$ |
| --- | --- | --- | --- |
| 1. high | $(\rho^* = 0.6)$ | 1-a | (0.2, 0.05, 0.05, 3.0, 3.0) |
|  |  | 1-b | (2.0, 0.7, 0.1, 2.5, 2.5) |
| 2. moderate | $(\rho^* = 0.3)$ | 2-a | (1.0, 1.0, 1.0, 1.5, 1.5) |
|  |  | 2-b | (3.0, 2.0, 1.0, 1.5, 0.5) |
| 3. low | $(\rho^* = 0.1)$ | 3-a | (0.2, 0.3, 3.0, 2.0, 1.5) |
|  |  | 3-b | (0.5, 2.0, 2.0, 0.5, 3.0) |
| 4. very low | $(\rho^* = 0.01)$ | 4-a | (0.01, 0.1, 1.0, 0.5, 0.5) |
|  |  | 4-b | (0.05, 2.0, 3.0, 3.0, 0.5) |
| zero-inflation | | | $(\pi_1, \pi_2, \pi_3, \pi_4)$ |
| i. low |  |  | (0.7, 0.1, 0.1, 0.1) |
| ii. moderate-balanced |  |  | (0.5, 0.15, 0.15, 0.2) |
| III. moderate-unbalanced |  |  | (0.5, 0.1, 0.3, 0.1) |
| iv. high-balanced |  |  | (0.2, 0.2, 0.2, 0.4) |
| v. high-unbalanced |  |  | (0.2, 0.1, 0.4, 0.3) |

For each *k* of *n*_sim_ simulation replicates, we got an estimate 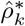 of the parameter *ρ*^*^, the standard error estimate *se*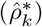, and the logit-transformed 95% confidence interval (i.e., 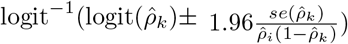). Then for each set of parameters, the following three quantities were calculated:

- the average estimated standard error (SE, 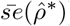)
- the standard deviation of the parameter estimates (SD, 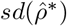)
- the empirical coverage probability (CP, 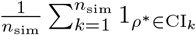, where CI_*k*_ is the logit-transformed 95% confidence interval for the *k*th replicate).

The simulation results are provided in Figures 5 and 6. First, the mean parameter estimates are close, or getting closer as sample size grows, to their true parameter values for each of the 40 scenarios. For most of the 40 parameter sets, CP was close to 0.95, and for those not close, CP gets closer to 0.95 with increasing sample size. In the same context, the average estimated standard error (SE) was close to the standard deviation of the parameter estimates (SD) especially when the sample size was large. However, when the underlying correlation was close to zero (i.e., 0.01 in our example), standard error estimation did not perform as well in terms of both CP and closeness of SE to SD. The parameter value being near the boundary may be responsible for the poorer performance. Also, Scenarios iv and v have higher SE and SD than the others. One possible explanation to this is that the effective sample size for those high zero-inflation scenarios is smaller than the other scenarios.

**Fig. 6.**
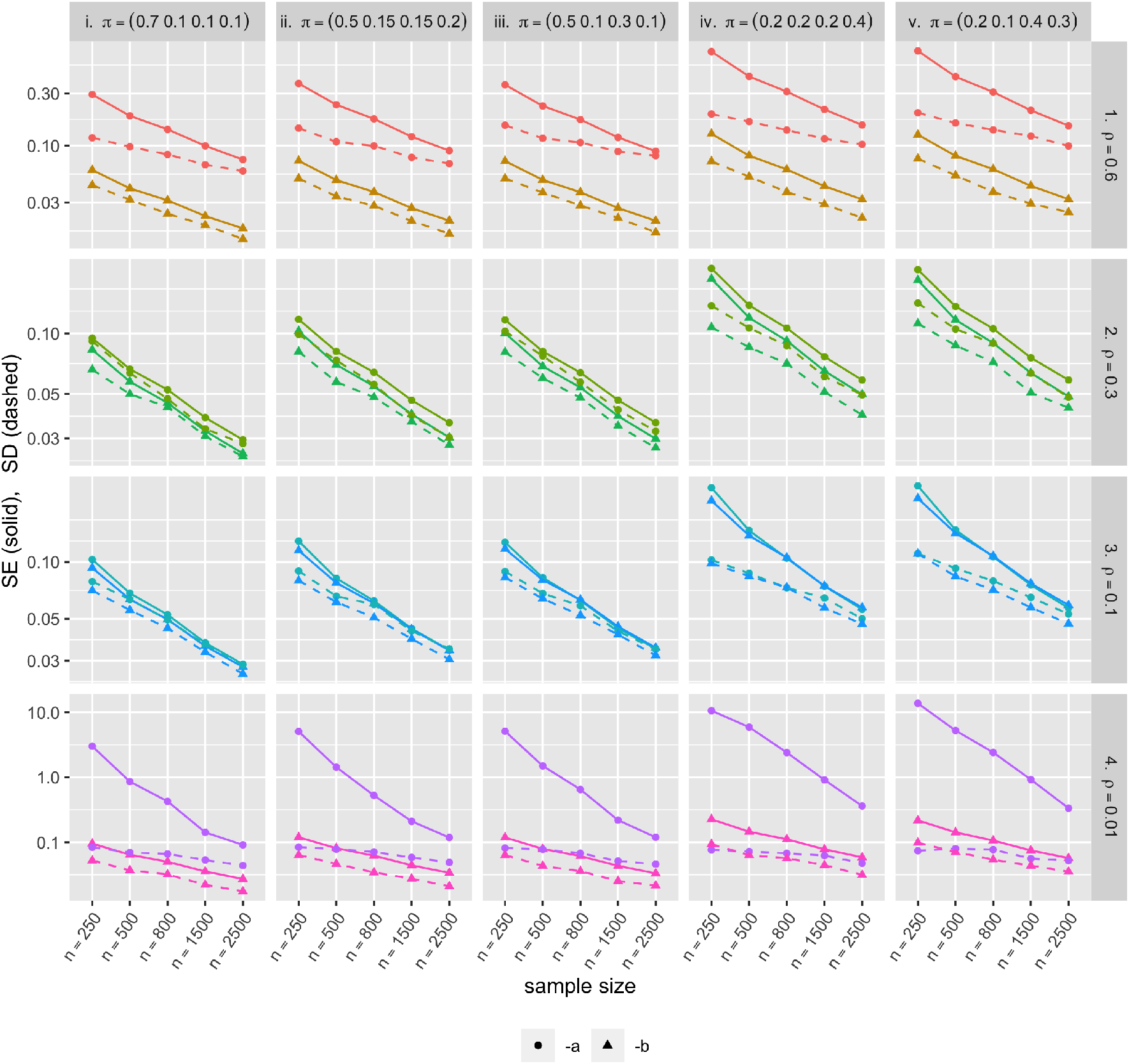
Standard error (SE, solid lines) and standard deviation (SD, dashed lines) of the BZINB-based underlying correlation estimates. Each color represents distinct simulation scenarios.

## 5. Modeling the incidence of dental caries on two surfaces using the BZINB model

In the dental caries clinical trial (X-ACT) study (Ritter *and others*, 2013), 647 participants (ages 21–80 years) were randomized to receive Xylitols versus inactive mints with 50% chance. The number of caries 36 months after treatment initiation was recorded for three types of surfaces: proximal, occlusal, and smooth surfaces with the proportion of no-caries for each type being 23.8%, 48.3%, and 22.6%, respectively. Figure 7 compares the empirical distribution with the model-estimates and provides the zero-inflation test results for each surface-type data. According to the test, the number of caries on the occlusal-surface, despite its high zero proportion, is not zero-inflated (*p* = 1.00), while that of the smooth-surface caries (no-caries for 23% of the sample) is significantly zero-inflated (*p* = 0.0025). The NB model does not seem to have a good fit to the proximal type, although the zero-inflation test statistic is not statistically significant at 5%. Thus, a BZINB model is suggested if the joint distribution between smooth-surface caries and one of the others is of interest.

**Fig. 7.**
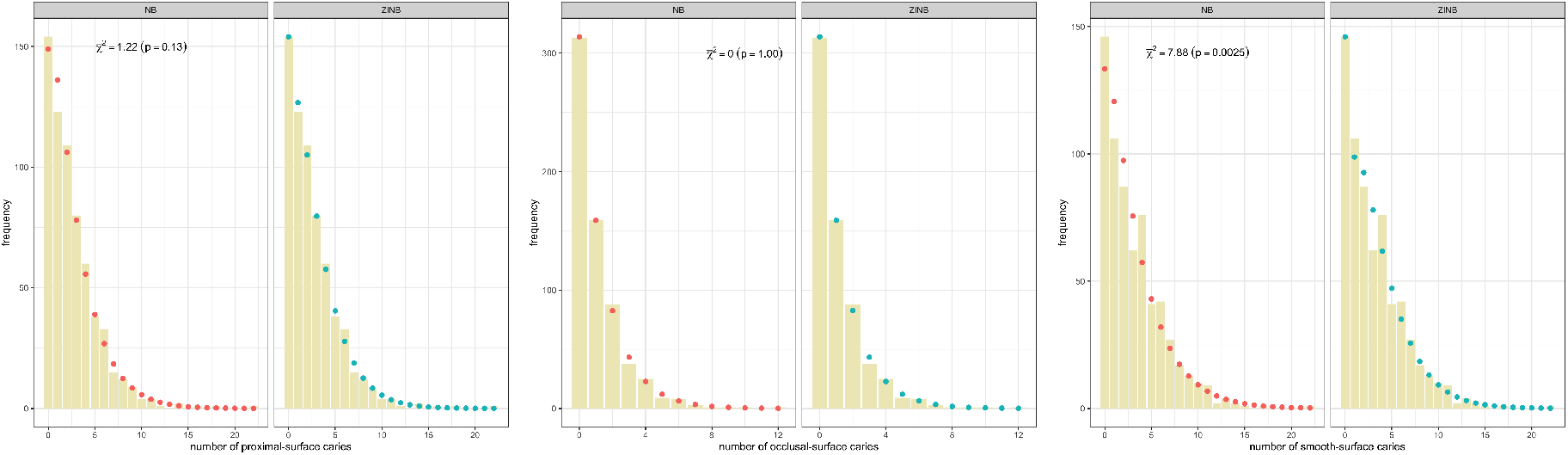
The empirical distribution of number of caries (bars). Proximal-, occlusal-, and smooth-surfaces for each of the three panels with the fitted NB distribution (red dots, LEFT), the fitted ZINB distribution (blue dots, RIGHT), and the zero-inflation test statistics 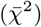 and their p-values.

We focus our analyses on the distribution of proximal- and smooth-surface caries, the counts of which are denoted as *Y*_1_ and *Y*_2_, respectively. The following two approaches shed light on the effectiveness and the efficiency of the bivariate model. First, we investigate the difference in the joint distribution of the caries counts on both surfaces between the intervention (Xylitol) and control (non-Xylitol) groups which is not obtainable from univariate models. In the second analysis, we illustrate how bivariate models could be more efficient than univariate models in testing marginal mean differences. The BZINB model parameter estimates for each group are provided in Table 2 and, together with their covariance estimates, the analyses of the joint dichotomized distribution and marginal mean tests are derived.

**Table 2.** The BZINB parameter estimates and their standard errors for the Xylitol and non-Xylitol groups

| parameter | non-Xylitol | Xylitol |
| --- | --- | --- |
| $(\alpha, \beta)$ | (1.51, 0.28, 0.008, 1.58, 2.34) | (1.56, 0.001, 0.14, 1.65, 1.86) |
| $SE(\hat{\alpha}, \hat{\beta})$ | (0.28, 0.18, 0.22, 0.28, 0.43) | (0.33, 0.21, 0.21, 0.25, 0.32) |
| $(\pi)$ | (0.936, 0.007, < 0.001, 0.057) | (0.921, 0.035, 0.009, 0.035) |
| $SE(\hat{\pi})$ | (0.04, 0.02, 0.02, 0.02) | (0.05, 0.03, 0.03, 0.03) |

In the first analysis, the joint distribution of the dichotomized caries statuses, or the prevalence (Pr(*Y*_1_ = 0, *Y*_2_ = 0), Pr(*Y*_1_ *>* 0, *Y*_2_ = 0), Pr(*Y*_1_ = 0, *Y*_2_ *>* 0)), is estimated for each group based on the BZINB model. See Table 3 for the estimates. The detailed formulae are provided in Web Appendix C1 of the Supplementary Materials. The BZINB model-based estimates are fairly close to the empirical distribution (Web Table 1 in Supplementary Materials) and provide, at the same time, a more stable and systematic estimation of the true distribution than the empirical distribution estimators.

**Table 3.**
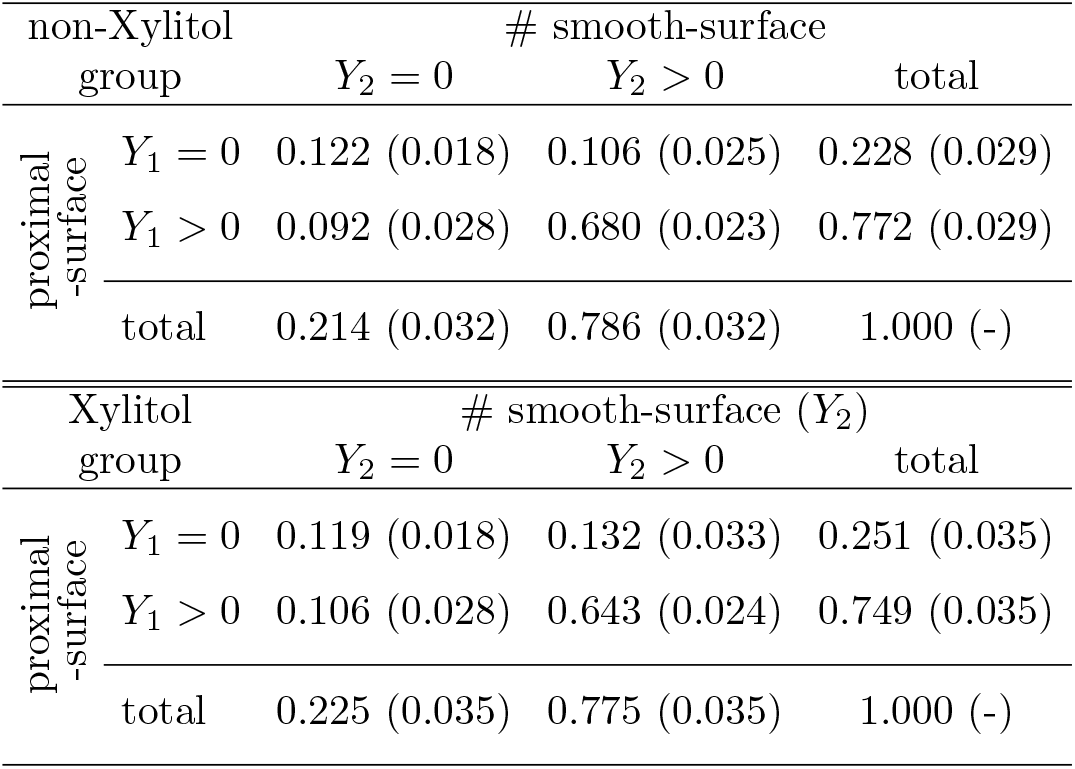
The BZINB-estimated joint probability (and its standard error) of dichotomized caries incidence on smooth- and proximal-surfaces for non-Xylitol (top) and Xylitol (bottom) groups. The estimated probabilities are given by the plug-in estimates of (2.4) for each group. See Web Appendix C2 for more details.

For both proximal and smooth surfaces, an increase in the marginal proportion of caries-free participants is observed in the treatment group. However, interestingly, the proportion of either caries or caries-free for both surfaces jointly has decreased. Provided that there was virtually no change in caries-free-for-both-surfaces, this perhaps implies a shift from caries-for-both-surfaces to caries-for-one-surface. Approximately 3.7%p, where “%p” refers to percentage points, of the caries-for-both group have been transferred to caries-for-smooth-surface-only (2.6%p) or caries-for-proximal-surface-only (1.4%p) groups, with 0.3%p difference is due to decrease in caries-free-for-both group.

In the second analysis, we compare the overall mean caries counts between the Xylitol and control groups for the two outcomes because, in clinical trials, the overall means are typically of interest as opposed to the latent class means. Nonetheless, two-part models for counts provide the structure to test the difference in the marginal means, *E*[*Y*_*j*_|non-Xylitol] − *E*[*Y*_*j*_|Xylitol] for *j* = 1, 2, using both the uni- and bivariate ZINB models. Since the Xylitol intervention is not expected to worsen the incidence, directional tests are used. The derivation of the statistics is relegated to Web Appendix C3 of the Supplementary Materials.

It can be seen from Table 4 that while the means are not very different across the models being used, the use of the bivariate model appears to enhance the power of the test owing to the increased precision of the marginal means estimates. This result is as expected, because in univariate models, the information contained in the other variable not being modeled is not utilized, while bivariate models can leverage the information of the other variable implied by the underlying structure. The standard errors of the BZINB-based estimators (in parentheses) and the p-values of the BZINB-based tests are overall smaller than those of the ZINB-based ones. Most notably, the p-value of the univariate test for smooth surfaces is 0.051 under the BZINB model, compared to 0.106 for the ZINB model.

**Table 4.**
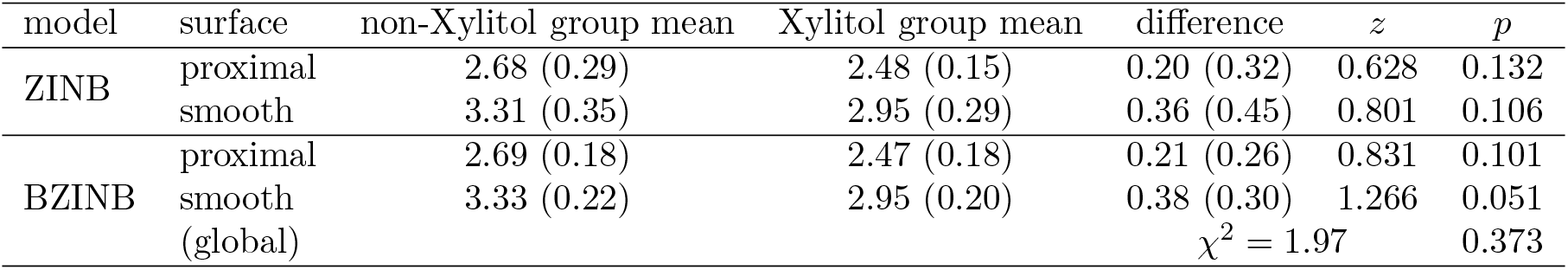
The univariate and bivariate model-based estimates (and the standard errors in parentheses) of the group means and the group differences. The *p*-values are given by the directional z-tests. The global mean difference test is done with the chi-square distribution of two degrees of freedom as the reference.

| model | surface | non-Xylitol group mean | Xylitol group mean | difference | $z$ | $p$ |
| --- | --- | --- | --- | --- | --- | --- |
| ZINB | proximal | 2.68 (0.29) | 2.48 (0.15) | 0.20 (0.32) | 0.628 | 0.132 |
|  | smooth | 3.31 (0.35) | 2.95 (0.29) | 0.36 (0.45) | 0.801 | 0.106 |
| BZINB | proximal | 2.69 (0.18) | 2.47 (0.18) | 0.21 (0.26) | 0.831 | 0.101 |
|  | smooth | 3.33 (0.22) | 2.95 (0.20) | 0.38 (0.30) | 1.266 | 0.051 |
| | (global) | | | $\chi^2 = 1.97$ | | 0.373 |

A further advantage of bivariate models is the use of global tests where inference is made by testing against the null hypothesis that the marginal mean parameters of the two groups are the same for all surfaces and rejecting the null if the marginal means differ between groups for at least one of the surface types. In the Xylitol trial, the BZINB-based global test of differences is not statistically significant (p=0.373), therefore we fail to reject the null hypothesis of no difference of average proximal and smooth surface caries counts between Xylitol and non-Xylitol mints.

## 6. Discussion

This article proposes a richly parametrized BZINB model that provides a full specification for the distribution of two correlated, overdispersed and zero-inflated, count random variables. It models bivariate count data with high flexibility by having eight free parameters and at the same time with simple latent variable interpretations. The hierarchical nature of the framework allows the use of nested models, such as BNB and BZIP models, and makes the model highly versatile and applicable to various contexts. In the scRNA-seq settings, by decomposing two sources of zeros, the distribution of counts without dropouts is recovered and the dependence is measured accordingly. In a second example with a totally different perspective on the meaning and utility of modeling “excess zeros” than the first example, the joint pattern of two types of dental caries was examined using the BZINB model in the Xylitol mints clinical trial. In particular, the BZINB model applied to bivariate caries counts from the Xylitol study enabled estimation of marginal parameters of common interest in clinical trials including the joint probabilities for the presence versus absence of any caries and overall mean caries counts for each surface type.

The BZINB model proposed in this article assumes an independent and identically distributed random bivarate sample of zero-inflated counts. Future work could generalize this homogeneous mean model to allow for covariance analysis or joint conditional mean analysis by introducing the generalized linear model framework. As in the univariate ZINB regression, the latent count variables (i.e., *X*_1_ and *X*_2_) can be modeled using linear predictors with some link function. For example, in the Xylitol study, the treatment indicator or baseline factors could be included in the linear predictors. Nonetheless, we are easily able to construct z-tests based on separate BZINB models for the two treatment groups in the Xylitol study because the groups were statistically independent. In a similar way, tests of treatment efficacy could be conducted for independent subgroups according to baseline factors, although often such tests have limited power.

There is a growing amount of literature that many scRNAseq data are not zero-inflated, and dropout events are primarily caused by PCR amplification that could be removed by the unique molecular identifiers (UMI) technique (Vieth *and others*, 2017; Townes *and others*, 2019; Svensson, 2020). While a good amount of comfort is available that there is no zero-inflation in the data for the droplet-based data such as 10X that uses UMI quantification, there is still a need to address dropouts in other platform-based scRNA-seq data as well as single cell proteomics and metatranscriptomics data. As we have observed the presence of zero-inflation in Section 4, zero-inflated models such as BZINB are needed in the example dataset.

Our model can be applied to other settings where there is a belief in two sources of zeros such as frailty, e.g., the first source corresponds to a cohort of people who are not susceptible to disease and will always have a zero count; the other source are random zeros among susceptible individuals. In this case, the dependence measure proposed in equation (2.9) applies to the bivariate outcome among the latent class of individuals that are susceptible to disease.

The proper use of the BZINB model depends on researchers’ understanding of how zeros were generated in the data. For example, if the expressed mRNAs are captured and sequenced without dropouts with a certain platform, the observed zeros in the resulting data would represent genes with no expression. In these settings where the excess zeros are not caused by dropout, the overall mean count and the proportion of subjects with positive counts have meaningful interpretations that may be directly modeled by marginalized ZINB (Preisser *and others*, 2016) and hurdle models (Mullahy, 1986), respectively. Directly modeling the observed pairs of counts, i.e., (*Y*_1_, *Y*_2_), using such models extended to bivariate counts could be beneficial, including for clinical trials such as the Xylitol study where marginal means are of interest and zero-inflated models are viewed as a convenient mechanism to account for many zeros (Mwalili *and others*, 2008). These scenarios underscore that any model, including BZINB, may not be ideal for all purposes, and that the statistical model for zero-inflated counts should be chosen to match the research question (Preisser *and others*, 2017).

In the BZINB model, allowing only positive *ρ*^*^ can be regarded as a limitation. One justification for the BZINB model is that the negative correlation of count data is not so prevalent in reality. For example, in genomics data, there are some genes that suppress other genes from being expressed, however, such genes either are relatively rare or have weak negative correlation with other genes. On the other hand, when we believe that the zeros are mostly not induced by dropout events, we can consider using the overall correlation (*ρ*(*Y*_1_, *Y*_2_)) which allows for negative correlation, instead of *ρ*^*^(*Y*_1_, *Y*_2_).

Finally, the BZINB model can also be generalized to a multivariate zero-inflated negative binomial model. This model may have an exponentially increasing number of latent variables or parameters as the dimension gets large. Though the lack of parsimony may make the multivariate model look less attractive, the idea can be very practically used in simulating multivariate zero-inflated count data and potentially in statistical analysis based on Bayesian models. For instance, a genomic count data with large amount of zeros can be mimicked by a set of latent random layers along with the generalized linear model framework. In dental caries clinical trials, a trivariate ZINB model could analyze caries counts from three tooth surface types simultaneously with a single global test avoiding the need for multiplicity adjustments to control family-wise Type I error.

## 7. Software

An `R` package `bzinb` estimating BZINB parameters is available on CRAN. The R code for the mouse paneth and the dental caries data analyses and simulations is available on https://github.com/Hunyong/BZINB_analysis.

## 8. Supplementary Material

The reader is referred to the on-line Supplementary Materials for the standard error calculation, details of the EM algorithm, additional details of the Xylitol experiment data analyses, and Web Figures. Supplementary material is available

## Acknowledgments

This work was supported by grant from the National Institutes of Health, National Institute of Dental and Craniofacial Research, R03-DE028983, and University of North Carolina Computational Medicine Program Award 2020. The authors thank Scott Magness and Joshua Starmer for providing the mouse paneth scRNA seq data and Michael I. Love for discussion of the implications of the dropouts.

