## Supplementary Materials for "A bivariate zero-inflated negative binomial model and its applications to biomedical settings"

*bioRxiv* (2021), **0**, 0, *pp.* 1–13

### Supplementary Materials for "A bivariate zero-inflated negative binomial model for identifying underlying dependence with application to single cell RNA sequencing data" by Hunyong Cho, Chuwen Liu, John S. Preisser, and Di Wu

Hunyong Cho, Chuwen Liu, John S. Preisser, and Di Wu

© The Author 2021. Published by Oxford University Press. All rights reserved. For permissions, please

#### Web Appendix A. EM algorithm

*The components of expected log-likelihood*

The expectation of the log-likelihood comprises the folowing terms with subscripts $i$ being suppressed:

$$E(R_0|Y_1,Y_2;\theta) = \alpha_0\beta_1 \frac{f_{BZINB}(Y_1,Y_2;\alpha_0+1,\alpha_1,\alpha_2,\beta,\pi)}{f_{BZINB}(Y_1,Y_2;\alpha_0,\alpha_1,\alpha_2,\beta,\pi)},$$

$$E(R_1|Y_1,Y_2;\theta) = \alpha_1\beta_1 \frac{f_{BZINB}(Y_1,Y_2;\alpha_0,\alpha_1+1,\alpha_2,\beta,\pi)}{f_{BZINB}(Y_1,Y_2;\alpha_0,\alpha_1,\alpha_2,\beta,\pi)},$$

$$E(R_2|Y_1,Y_2;\theta) = \alpha_2\beta_1 \frac{f_{BZINB}(Y_1,Y_2;\alpha_0,\alpha_1,\alpha_2+1,\beta,\pi)}{f_{BZINB}(Y_1,Y_2;\alpha_0,\alpha_1,\alpha_2,\beta,\pi)},$$

$$\begin{aligned} E(\log(R_0)|Y_1,Y_2;\theta) = & \frac{1}{f_{BZINB}(Y_1,Y_2)}\times \\ & \Bigg[\sum_{k,m} H_0(k,m;Y_1,Y_2,\alpha,\beta)\{\psi(Y_1+Y_2-k-m+\alpha_0)+\log(\frac{\beta_1}{1+\beta_1+\beta_2})\}\pi_1+ \\ & \sum_{k=0}^{Y_1} H_1(k;Y_1,\alpha,\beta)\{\psi(Y_1-k+\alpha_0)+\log(\frac{\beta_1}{1+\beta_1})\}\pi_2\zeta(Y_2)+ \\ & \sum_{m=0}^{Y_2} H_2(m;Y_2,\alpha,\beta)\{\psi(Y_2-m+\alpha_0)+\log(\frac{\beta_1}{1+\beta_2})\}\pi_3\zeta(Y_1)+ \\ & \{\psi(\alpha_0)+\log(\beta_1)\}\pi_4\zeta(Y_1+Y_2)\Bigg], \end{aligned}$$

$$\begin{aligned} E(\log(R_1)|Y_1,Y_2;\theta) = & \frac{1}{f_{BZINB}(Y_1,Y_2)}\times \\ & \Bigg[\sum_{k,m} H_0(k,m;Y_1,Y_2,\alpha,\beta)\{\psi(k+\alpha_1)+\log(\frac{\beta_1}{1+\beta_1})\}\pi_1+ \\ & \sum_{k=0}^{Y_1} H_1(k;Y_1,\alpha,\beta)\{\psi(k+\alpha_1)+\log(\frac{\beta_1}{1+\beta_1})\}\pi_2\zeta(Y_2)+ \\ & \sum_{m=0}^{Y_2} H_2(m;Y_2,\alpha,\beta)\{\psi(\alpha_1)+\log(\beta_1)\}\pi_3\zeta(Y_1)+ \\ & \{\psi(\alpha_1)+\log(\beta_1)\}\pi_4\zeta(Y_1+Y_2)\Bigg], \end{aligned}$$

$$\begin{aligned} E(\log(R_2)|Y_1,Y_2;\theta) = & \frac{1}{f_{BZINB}(Y_1,Y_2)}\times \\ & \Bigg[\sum_{k,m} H_0(k,m;Y_1,Y_2,\alpha,\beta)\{\psi(m+\alpha_2)+\log(\frac{\beta_1}{1+\beta_2})\}\pi_1+ \\ & \sum_{k=0}^{Y_1} H_1(k;Y_1,\alpha,\beta)\{\psi(\alpha_2)+\log(\beta_1)\}\pi_2\zeta(Y_2)+ \\ & \sum_{m=0}^{Y_2} H_2(m;Y_2,\alpha,\beta)\{\psi(m+\alpha_2)+\log(\frac{\beta_1}{1+\beta_2})\}\pi_3\zeta(Y_1)+ \\ & \{\psi(\alpha_2)+\log(\beta_1)\}\pi_4\zeta(Y_1+Y_2)\Bigg], \end{aligned}$$

$$
\begin{aligned}
E(E_1|Y_1, Y_2; \theta) &= \frac{f_{BNB}(Y_1, Y_2; \alpha, \beta)\pi_1}{f_{BZINB}(Y_1, Y_2; \theta)}, \\
E(E_2|Y_1, Y_2; \theta) &= \frac{f_{NB}(Y_1; \alpha_0 + \alpha_1, \frac{\beta_1}{\beta_1+1})\pi_2\zeta(Y_2)}{f_{BZINB}(Y_1, Y_2; \theta)}, \\
E(E_3|Y_1, Y_2; \theta) &= \frac{f_{NB}(Y_2; \alpha_0 + \alpha_2, \frac{\beta_2}{\beta_2+1})\pi_3\zeta(Y_1)}{f_{BZINB}(Y_1, Y_2; \theta)}, \\
E(E_4|Y_1, Y_2; \theta) &= \frac{\pi_4\zeta(Y_1 + Y_2)}{f_{BZINB}(Y_1, Y_2; \theta)}, \qquad \text{and} \\
E(X_2|Y_1, Y_2; \theta) &= Y_2 + \frac{(\alpha_0 + \alpha_2)\beta_2}{f_{BZINB}(Y_1, Y_2; \theta)} \times \\
&\quad \left[ f_{NB}(Y_1; \alpha_0 + \alpha_1, \frac{\beta_1}{\beta_1 + 1})\pi_2\zeta(Y_2) + \pi_4\zeta(Y_1 + Y_2) \right],
\end{aligned}
$$

where

$$
\begin{aligned}
H_0(k, m; Y_1, Y_2, \alpha, \beta) &:= \binom{\alpha_0 + Y_1 + Y_2 - k - m - 1}{\alpha_0 + Y_2 - m - 1} \binom{\alpha_0 + Y_2 - m - 1}{\alpha_0 - 1} \times \\
&\quad \binom{\alpha_1 + k - 1}{\alpha_1 - 1} \binom{\alpha_2 + m - 1}{\alpha_2 - 1} \frac{\beta_1^{Y_1}\beta_2^{Y_2}(\beta_1 + \beta_2 + 1)^{k+m-Y_1-Y_2-\alpha_0}}{(\beta_1 + 1)^{k+\alpha_1}(\beta_2 + 1)^{m+\alpha_2}}, \\
H_1(k; Y_1, \alpha, \beta) &:= \binom{\alpha_0 + Y_1 - k - 1}{\alpha_0 - 1} \binom{\alpha_1 + k - 1}{\alpha_1 - 1} \frac{\beta_1^{Y_1}}{(\beta_1 + 1)^{Y_1+\alpha_0+\alpha_1}}, \\
H_2(m; Y_2, \alpha, \beta) &:= \binom{\alpha_0 + Y_2 - m - 1}{\alpha_0 - 1} \binom{\alpha_2 + m - 1}{\alpha_2 - 1} \frac{\beta_2^{Y_2}}{(\beta_2 + 1)^{Y_2+\alpha_0+\alpha_2}},
\end{aligned}
$$

$\theta \equiv (\alpha^\top, \beta^\top, \pi^\top)^\top$, $\psi(\cdot)$ is the digamma function, and the parameters in density functions are written either in scalar, vector, or combination of both as needed without confusion.

#### Web Appendix B. Standard error formula

Recall the density of BZINB is given as,

$$
\begin{aligned}
&f_{BZINB}(y_1, y_2; \theta) \\
&= \pi_1 f_{BNB}(y_1, y_2; \theta) + \pi_2 f_{NB}(y_1; \alpha_0 + \alpha_1, \frac{1}{\beta_1 + 1})\zeta(y_2) \\
&\quad + \pi_3 f_{NB}(y_2; \alpha_0 + \alpha_2, \frac{1}{\beta_2 + 1})\zeta(y_1) + (1 - \pi_1 - \pi_2 - \pi_3)\zeta(y_1 + y_2),
\end{aligned}
$$

where $\theta$ here is redefined using free parameters only as $(\alpha^\top, \beta^\top, \pi_1, \pi_2, \pi_3)^\top$.

Then the observed information is given as,

$$
\begin{aligned}
I_{obs}(\hat\theta) &= -\partial_\theta^2 l|_{\theta=\hat\theta} \\
&= -\sum_i^n \partial_\theta^2 \log f_{BZINB}(y_{1,i}, y_{2,i};\theta)|_{\theta=\hat\theta} \\
&= -\sum_i^n \frac{\{\partial_\theta^2 f_{BZINB}(y_{1,i}, y_{2,i};\theta)\}\{f_{BZINB}(y_{1,i}, y_{2,i};\theta)\} - \{\partial_\theta f_{BZINB}(y_{1,i}, y_{2,i};\theta)\}^{\otimes 2}}{\{f_{BZINB}(y_{1,i}, y_{2,i};\theta)\}^2}|_{\theta=\hat\theta},
\end{aligned}
$$

where $a^{\otimes 2} = aa^\top$ and $\partial_\theta^2 l = \partial_\theta \partial_{\theta^\top} l$.

Then the large sample standard error estimate of the MLE $\hat\theta$ of $\theta$ is $diag(I_{obs}(\hat\theta)^{-1})^{\frac{1}{2}}$, and that of the MLE $\hat\rho^*$ of $\rho^*$ is $\left[(\nabla g(\hat\theta))^\top I_{obs}(\hat\theta)^{-1} \nabla g(\hat\theta)\right]^{1/2}$, where $\nabla g(\hat\theta) := \partial_\theta \rho^*|_{\hat\theta}$ is as follows:

$$
\begin{aligned}
\nabla g(\hat\theta) = \hat\rho^{TRUE} \Bigg( & \left\{ \frac{1}{\hat\alpha_0} - \frac{1}{2(\hat\alpha_0 + \hat\alpha_1)} - \frac{1}{2(\hat\alpha_0 + \hat\alpha_2)} \right\}, \frac{-1}{2(\hat\alpha_0 + \hat\alpha_1)}, \frac{-1}{2(\hat\alpha_0 + \hat\alpha_2)}, \\
& \frac{1}{2\hat\beta_1(\hat\beta_1 + 1)}, \frac{1}{2\hat\beta_2(\hat\beta_2 + 1)}, 0, 0, 0 \Bigg)^\top .
\end{aligned}
$$

The observed information, however, can be approximated by the empirical observed information (e.g. Meilijson (1989)) which incurs only the first derivatives of the individual log-likelihood:

$$
I_e(\hat\theta) = \sum_{i=1}^n s(y_{1,i}, y_{2,i};\hat\theta) s(y_{1,i}, y_{2,i};\hat\theta)^\top,
$$

where

$$
\begin{aligned}
s(y_{1,i}, y_{2,i};\theta) &= \partial_\theta \log f_{BZINB}(y_{1,i}, y_{2,i};\theta) \\
&= \frac{\partial_\theta f_{BZINB}(y_{1,i}, y_{2,i};\theta)}{f_{BZINB}(y_{1,i}, y_{2,i};\theta)},
\end{aligned}
$$

$$
\partial_\theta f_{BZINB}(y_1, y_2;\theta) = \begin{pmatrix}
\pi_1 D_1 + \pi_2 \zeta(y_2) D_6 + \pi_3 \zeta(y_1) D_8 \\
\pi_1 D_2 + \pi_2 \zeta(y_2) D_6 \\
\pi_1 D_3 + \pi_3 \zeta(y_1) D_8 \\
\pi_1 D_4 + \pi_2 \zeta(y_2) D_7 \\
\pi_1 D_5 + \pi_3 \zeta(y_1) D_9 \\
f_{BNB}(y_1, y_2;\alpha, \beta) - \zeta(y_1 + y_2) \\
\zeta(y_2) f_{NB}(y_1;\alpha_0 + \alpha_1, \frac{1}{\beta_1 + 1}) - \zeta(y_1 + y_2) \\
\zeta(y_1) f_{NB}(y_2;\alpha_0 + \alpha_2, \frac{1}{\beta_2 + 1}) - \zeta(y_1 + y_2)
\end{pmatrix},
$$

$$
\begin{aligned}
D_1 &= \partial_{\alpha_0} f_{BNB}(y_1, y_2; \alpha, \beta) \\
&= \sum_{k=0}^{y_1} \sum_{m=0}^{y_2} \Big[ H_0(k, m; y_1, y_2, \alpha, \beta) \Big\{ \psi(\alpha_0 + y_1 + y_2 - k - m) - \psi(\alpha_0) - \log(\beta_1 + \beta_2 + 1) \Big\} \Big], \\
D_2 &= \partial_{\alpha_1} f_{BNB}(y_1, y_2; \alpha, \beta) \\
&= \sum_{k=0}^{y_1} \sum_{m=0}^{y_2} \Big[ H_0(k, m; y_1, y_2, \alpha, \beta) \{ \psi(\alpha_1 + k) - \psi(\alpha_1) - \log(\beta_1 + 1) \} \Big], \\
D_3 &= \partial_{\alpha_2} f_{BNB}(y_1, y_2; \alpha, \beta) \\
&= \sum_{k=0}^{y_1} \sum_{m=0}^{y_2} \Big[ H_0(k, m; y_1, y_2, \alpha, \beta) \{ \psi(\alpha_2 + m) - \psi(\alpha_2) - \log(\beta_2 + 1) \} \Big], \\
D_4 &= \partial_{\beta_1} f_{BNB}(y_1, y_2; \alpha, \beta) \\
&= \sum_{k=0}^{y_1} \sum_{m=0}^{y_2} \Big[ H_0(k, m; y_1, y_2, \alpha, \beta) \Big\{ \frac{y_1}{\beta_1} - \frac{\alpha_0 + y_1 + y_2 - k - m}{\beta_1 + \beta_2 + 1} - \frac{k + \alpha_1}{\beta_1 + 1} \Big\} \Big], \\
D_5 &= \partial_{\beta_2} f_{BNB}(y_1, y_2; \alpha, \beta) \\
&= \sum_{k=0}^{y_1} \sum_{m=0}^{y_2} \Big[ H_0(k, m; y_1, y_2, \alpha, \beta) \Big\{ \frac{y_2}{\beta_2} - \frac{\alpha_0 + y_1 + y_2 - k - m}{\beta_1 + \beta_2 + 1} - \frac{m + \alpha_2}{\beta_2 + 1} \Big\} \Big],
\end{aligned}
$$

$$
\begin{aligned}
D_6 &= \partial_{\alpha_j} f_{NB}(y_1; \alpha_0 + \alpha_1, \frac{1}{\beta_1 + 1}) \qquad\qquad j = 0, 1 \\
&= f_{NB}(y_1; \alpha_0 + \alpha_1, \frac{1}{\beta_1 + 1}) \Big[ \psi(\alpha_0 + \alpha_1 + y_1) - \psi(\alpha_0 + \alpha_1) - \log(\beta_1 + 1) \Big], \\
D_7 &= \partial_{\beta_1} f_{NB}(y_1; \alpha_0 + \alpha_1, \frac{1}{\beta_1 + 1}) \\
&= f_{NB}(y_1; \alpha_0 + \alpha_1, \frac{1}{\beta_1 + 1}) \Big[ \frac{y_1}{\beta_1} - \frac{\alpha_0 + \alpha_1 + y_1}{\beta_1 + 1} \Big], \\
D_8 &= \partial_{\alpha_j} f_{NB}(y_2; \alpha_0 + \alpha_2, \frac{1}{\beta_2 + 1}) \qquad\qquad j = 0, 2 \\
&= f_{NB}(y_2; \alpha_0 + \alpha_2, \frac{1}{\beta_2 + 1}) \Big[ \psi(\alpha_0 + \alpha_2 + y_2) - \psi(\alpha_0 + \alpha_2) - \log(\beta_2 + 1) \Big], \text{ and} \\
D_9 &= \partial_{\beta_2} f_{NB}(y_2; \alpha_0 + \alpha_2, \frac{1}{\beta_2 + 1}) \\
&= f_{NB}(y_2; \alpha_0 + \alpha_2, \frac{1}{\beta_2 + 1}) \Big[ \frac{y_2}{\beta_2} - \frac{\alpha_0 + \alpha_2 + y_2}{\beta_2 + 1} \Big].
\end{aligned}
$$

#### Web Appendix C. Additional details of the Xylitol experiment data analyses

##### *Web Appendix C1. The formulae for the joint density estimation*

The dichotomized joint probabilities are derived as follows, and the plug-in estimates are given by replacing the parameters with the corresponding parameter estimates.

$$
\begin{aligned}
\Pr(Y_1 = Y_2 = 0) &= \frac{\pi_1}{(\beta_1+1)^{\alpha_1}(\beta_2+1)^{\alpha_2}(\beta_1+\beta_2+1)^{\alpha_0}} + \frac{\pi_2}{(\beta_1+1)^{\alpha_0+\alpha_1}} + \frac{\pi_3}{(\beta_2+1)^{\alpha_0+\alpha_2}} + \pi_4, \qquad (0.1)\\
&=: (J_1) + (J_2) + (J_3) + (J_4)\\
\Pr(Y_1 = 0) &= \frac{\pi_1+\pi_2}{(\beta_1+1)^{\alpha_0+\alpha_1}} + (\pi_3+\pi_4), \qquad (0.2)\\
&=: (K_1) + (K_2)\\
\Pr(Y_2 = 0) &= \frac{\pi_1+\pi_3}{(\beta_2+1)^{\alpha_0+\alpha_2}} + (\pi_2+\pi_4), \qquad (0.3)\\
&=: (L_1) + (L_2)\\
\Pr(Y_2 > Y_1 = 0) &= (0.2) - (0.1),\\
&= \frac{\pi_1}{(\beta_1+1)^{\alpha_1}}\Big\{\frac{1}{(\beta_1+1)^{\alpha_2}} - \frac{1}{(\beta_1+\beta_2+1)^{\alpha_0}(\beta_2+1)^{\alpha_2}}\Big\} + \pi_3\Big\{1 - \frac{1}{(\beta_2+1)^{\alpha_0+\alpha_2}}\Big\} \qquad (0.4)\\
\Pr(Y_1 > Y_2 = 0) &= (0.3) - (0.1),\\
&= \frac{\pi_2}{(\beta_2+1)^{\alpha_2}}\Big\{\frac{1}{(\beta_2+1)^{\alpha_1}} - \frac{1}{(\beta_1+\beta_2+1)^{\alpha_0}(\beta_1+1)^{\alpha_1}}\Big\} + \pi_2\Big\{1 - \frac{1}{(\beta_1+1)^{\alpha_0+\alpha_1}}\Big\},\\
\Pr(Y_1 > 0, Y_2 > 0) &= 1 + (0.1) - (0.2) - (0.3)\\
&= \pi_1\Big\{1 + \frac{1}{(\beta_1+1)^{\alpha_1}(\beta_2+1)^{\alpha_2}(\beta_1+\beta_2+1)^{\alpha_0}} - \frac{1}{(\beta_1+1)^{\alpha_0+\alpha_1}} - \frac{1}{(\beta_2+1)^{\alpha_0+\alpha_2}}\Big\}. \qquad (0.5)
\end{aligned}
$$

Now, the variances of the above estimators are given by the following.

$$
\begin{aligned}
var(\widehat{\Pr}(Y_1 = Y_2 = 0)) &= \nabla g_A(\hat{\boldsymbol{\theta}})^\top var(\hat{\boldsymbol{\theta}}) \nabla g_A(\hat{\boldsymbol{\theta}}),\\
var(\widehat{\Pr}(Y_1 = 0)) &= \nabla g_B(\hat{\boldsymbol{\theta}})^\top var(\hat{\boldsymbol{\theta}}) \nabla g_B(\hat{\boldsymbol{\theta}}),\\
var(\widehat{\Pr}(Y_2 = 0)) &= \nabla g_C(\hat{\boldsymbol{\theta}})^\top var(\hat{\boldsymbol{\theta}}) \nabla g_C(\hat{\boldsymbol{\theta}}),\\
var(\widehat{\Pr}(Y_2 > Y_1 = 0)) &= \nabla g_D(\hat{\boldsymbol{\theta}})^\top var(\hat{\boldsymbol{\theta}}) \nabla g_D(\hat{\boldsymbol{\theta}}),\\
var(\widehat{\Pr}(Y_1 > Y_2 = 0)) &= \nabla g_E(\hat{\boldsymbol{\theta}})^\top var(\hat{\boldsymbol{\theta}}) \nabla g_E(\hat{\boldsymbol{\theta}}), \text{ and}\\
var(\widehat{\Pr}(Y_1, Y_2 > 0)) &= \nabla g_F(\hat{\boldsymbol{\theta}})^\top var(\hat{\boldsymbol{\theta}}) \nabla g_F(\hat{\boldsymbol{\theta}}),
\end{aligned}
$$

where

$$
\begin{aligned}
\nabla g_A(\boldsymbol{\theta}) &= \partial_{\boldsymbol{\theta}} \Pr(Y_1 = Y_2 = 0) \\
&= - \begin{pmatrix} (J_1 + J_2 + J_3) \log(\beta_1 + \beta_2 + 1) \\ (J_1 + J_2) \log(\beta_1 + 1) \\ (J_1 + J_3) \log(\beta_2 + 1) \\ J_1 \Big( \frac{\alpha_0}{\beta_1+\beta_2+1} + \frac{\alpha_1}{\beta_1+1} \Big) + J_2 \frac{\alpha_0+\alpha_1}{\beta_1+1} \\ J_1 \Big( \frac{\alpha_0}{\beta_1+\beta_2+1} + \frac{\alpha_1}{\beta_1+1} \Big) + J_3 \frac{\alpha_0+\alpha_2}{\beta_2+1} \\ 1 - J_1/\pi_1 \\ 1 - J_2/\pi_2 \\ 1 - J_3/\pi_3 \end{pmatrix}, \\
\nabla g_B(\boldsymbol{\theta}) &= \partial_{\boldsymbol{\theta}} \Pr(Y_1 = 0) \\
&= - \Big( K_1 \log(\beta_1 + 1), K_1 \log(\beta_1 + 1), 0, K_1 \frac{\alpha_0 + \alpha_1}{\beta_1 + 1}, 0, 1 - K_1/\pi_1, 1 - K_1/\pi_1, 0 \Big)^\top, \\
\nabla g_C(\boldsymbol{\theta}) &= \partial_{\boldsymbol{\theta}} \Pr(Y_2 = 0) \\
&= - \Big( L_1 \log(\beta_2 + 1), 0, L_1 \log(\beta_2 + 1), 0, L_1 \frac{\alpha_0 + \alpha_2}{\beta_2 + 1}, 1 - L_1/\pi_1, 0, 1 - L_1/\pi_1 \Big)^\top,
\end{aligned}
$$

$\nabla g_D = -\nabla g_A + \nabla g_B, \nabla g_E = -\nabla g_A + \nabla g_C$, and $\nabla g_F = -\nabla g_A + \nabla g_B + \nabla g_C$.

*Web Appendix C2. The empirical density of the dichotomized caries statuses*

Table 1. The empirical density of the dichotomized caries statuses for the proximal and smooth surfaces.

| non-Xylitol group | | # smooth-surface $Y_2 = 0$ | $Y_2 > 0$ | total |
|---|---|---|---|---|
| proximal -surface | $Y_1 = 0$ | 0.122 (0.013) | 0.113 (0.012) | 0.235 (0.017) |
| | $Y_1 > 0$ | 0.091 (0.011) | 0.674 (0.018) | 0.765 (0.017) |
| | total | 0.213 (0.017) | 0.787 (0.017) | 1.000 (-) |

| Xylitol group | | # smooth-surface ($Y_2$) $Y_2 = 0$ | $Y_2 > 0$ | total |
|---|---|---|---|---|
| proximal -surface ($Y_1$) | $Y_1 = 0$ | 0.119 (0.013) | 0.123 (0.013) | 0.242 (0.017) |
| | $Y_1 > 0$ | 0.119 (0.013) | 0.639 (0.019) | 0.758 (0.017) |
| | total | 0.238 (0.017) | 0.762 (0.017) | 1.000 (-) |

*Web Appendix C3. The formulae for the marginal mean difference tests*

Let $Y_j^G$ denote the number of caries on the $j$th surface for a person belonging to group $G$, $j = 1, 2, G = X, C$ and let $\mu_j^G = E[Y_j^G]$. We follow the same notational rule (super- and subscripts) for the elementary parameters $\boldsymbol{\theta}$. Then, for each $G$, we have from (2.6)

$$\begin{aligned}
(\mu_1^G, \mu_2^G) &= \big((\pi_1 + \pi_2)(\alpha_0 + \alpha_1)\beta_1, (\pi_1 + \pi_3)(\alpha_0 + \alpha_2)\beta_2\big) \\
cov((\hat{\mu}_1^G, \hat{\mu}_2^G)^\top) &= \nabla g_\mu^G(\theta)^\top cov(\hat{\boldsymbol{\theta}}) \nabla g_\mu^G(\theta) \\
&=: \begin{pmatrix} \sigma_{11}^G & \sigma_{12}^G \\ \sigma_{12}^G & \sigma_{22}^G \end{pmatrix}. \\
\nabla g_\mu^G(\theta) &= (\nabla g_{\mu_1}^G(\theta), \nabla g_{\mu_2}^G(\theta)) \\
\nabla g_{\mu_1}^G(\theta) &= ((\pi_1 + \pi_2)\beta_1, (\pi_1 + \pi_2)\beta_1, 0, (\pi_1 + \pi_2)(\alpha_0 + \alpha_1), 0, (\alpha_0 + \alpha_1)\beta_1, (\alpha_0 + \alpha_1)\beta_1, 0)^\top \\
\nabla g_{\mu_2}^G(\theta) &= ((\pi_1 + \pi_3)\beta_2, 0, (\pi_1 + \pi_3)\beta_2, 0, (\pi_1 + \pi_3)(\alpha_0 + \alpha_2), (\alpha_0 + \alpha_2)\beta_2, 0, (\alpha_0 + \alpha_2)\beta_2)^\top.
\end{aligned}$$

The corresponding estimates are obtained by plugging in $\hat{\theta}^G$ and are denoted with a hat.

The test statistics for the mean differences are given as

$$\frac{\hat{\mu}_j^C - \hat{\mu}_j^X}{\sqrt{\hat{\sigma}_{jj}^C + \hat{\sigma}_{jj}^X}}, j = 1, 2$$

and the chi-square statistic testing the global mean difference is given as

$$\hat{\Delta}^\top \hat{\Sigma}^{-1} \hat{\Delta}$$

with two degrees of freedom, where $\Delta = (\mu_1^C, \mu_2^C)^\top - (\mu_1^X, \mu_2^X)^\top$ and $\Sigma = \begin{pmatrix} \sigma_{11}^C & \sigma_{12}^C \\ \sigma_{12}^C & \sigma_{22}^C \end{pmatrix} + \begin{pmatrix} \sigma_{11}^X & \sigma_{12}^X \\ \sigma_{12}^X & \sigma_{22}^X \end{pmatrix}$.

#### Web Figures. Exhaustive outputs of real data examples

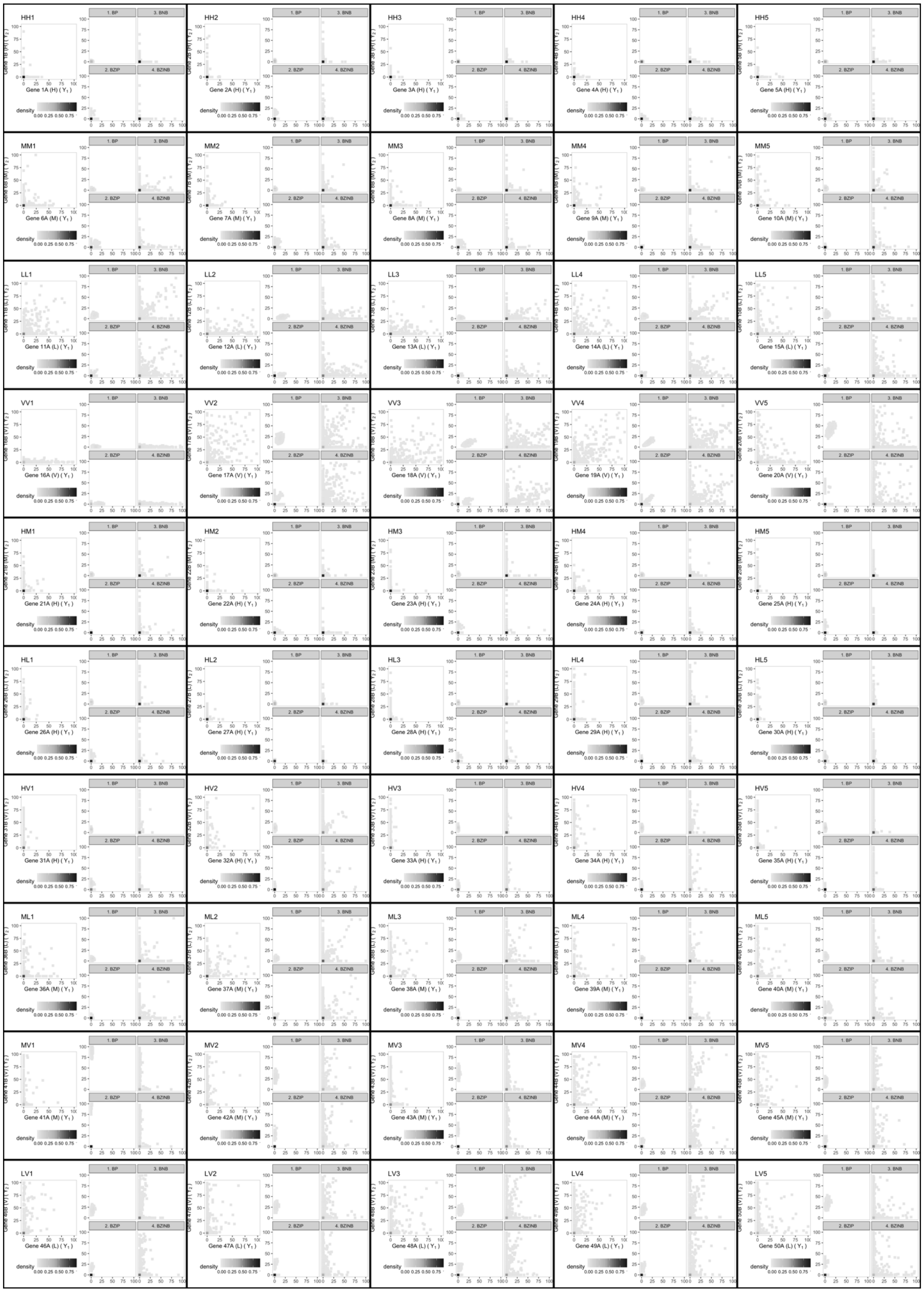

Fig. 1. The distribution of real and simulated mouse paneth RNA count data for all 50 pairs - the FIRST replicate. Each of the rows contains five pairs: From top to bottom, rows represent HH, MM, LL, VV, HM, HL, HV, ML, MV, and LV, in the order. Each pair has the real empirical distribution (LEFT), and the four model-based simulated empirical distributions (RIGHT).

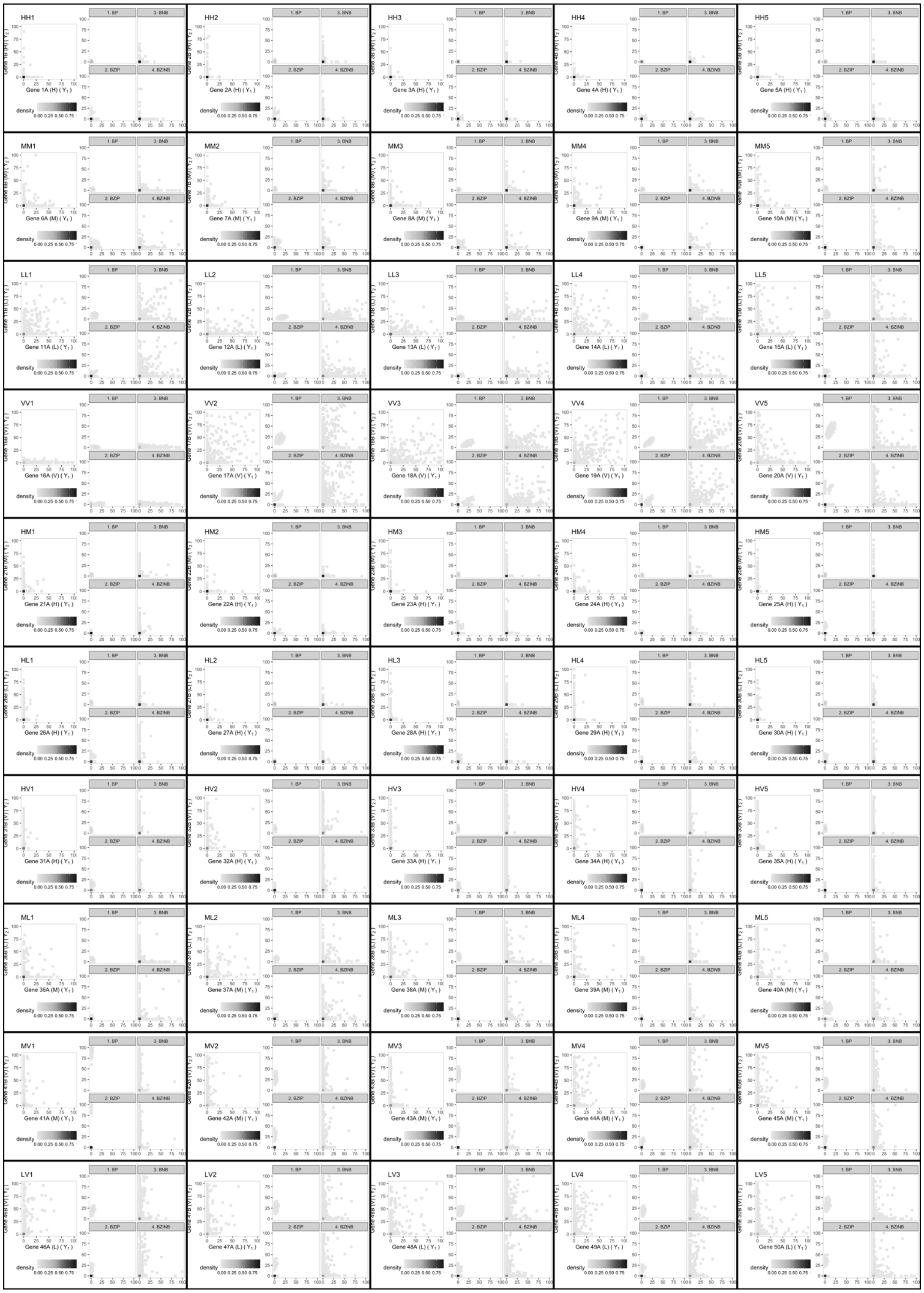

Fig. 2. The distribution of real and simulated mouse paneth RNA count data for all 50 pairs - the SECOND replicate. Each of the rows contains five pairs: From top to bottom, rows represent HH, MM, LL, VV, HM, HL, HV, ML, MV, and LV, in the order. Each pair has the real empirical distribution (LEFT), and the four model-based simulated empirical distributions (RIGHT).

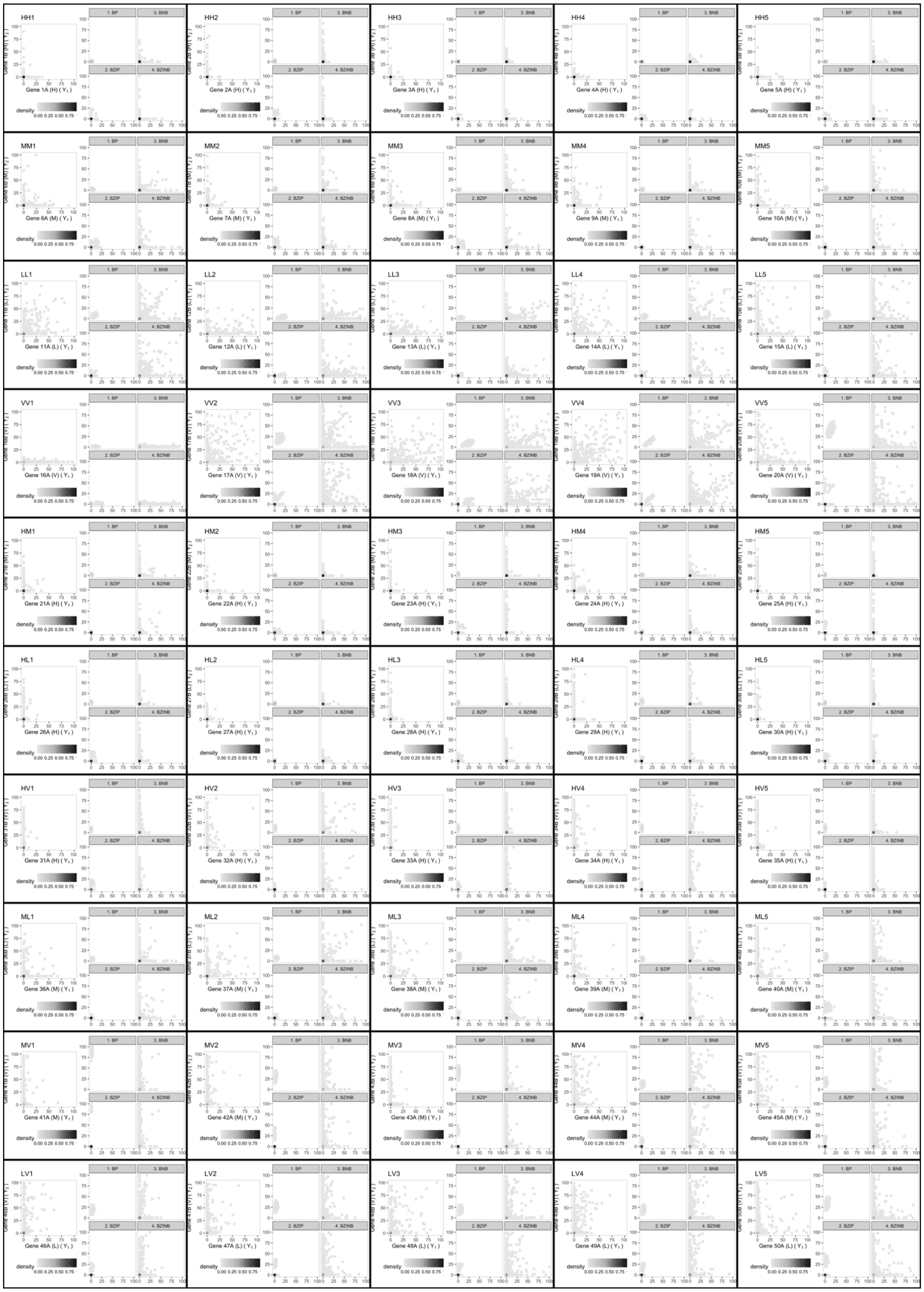


Fig. 3. The distribution of real and simulated mouse paneth RNA count data for all 50 pairs - the THIRD replicate. Each of the rows contains five pairs: From top to bottom, rows represent HH, MM, LL, VV, HM, HL, HV, ML, MV, and LV, in the order. Each pair has the real empirical distribution (LEFT), and the four model-based simulated empirical distributions (RIGHT).

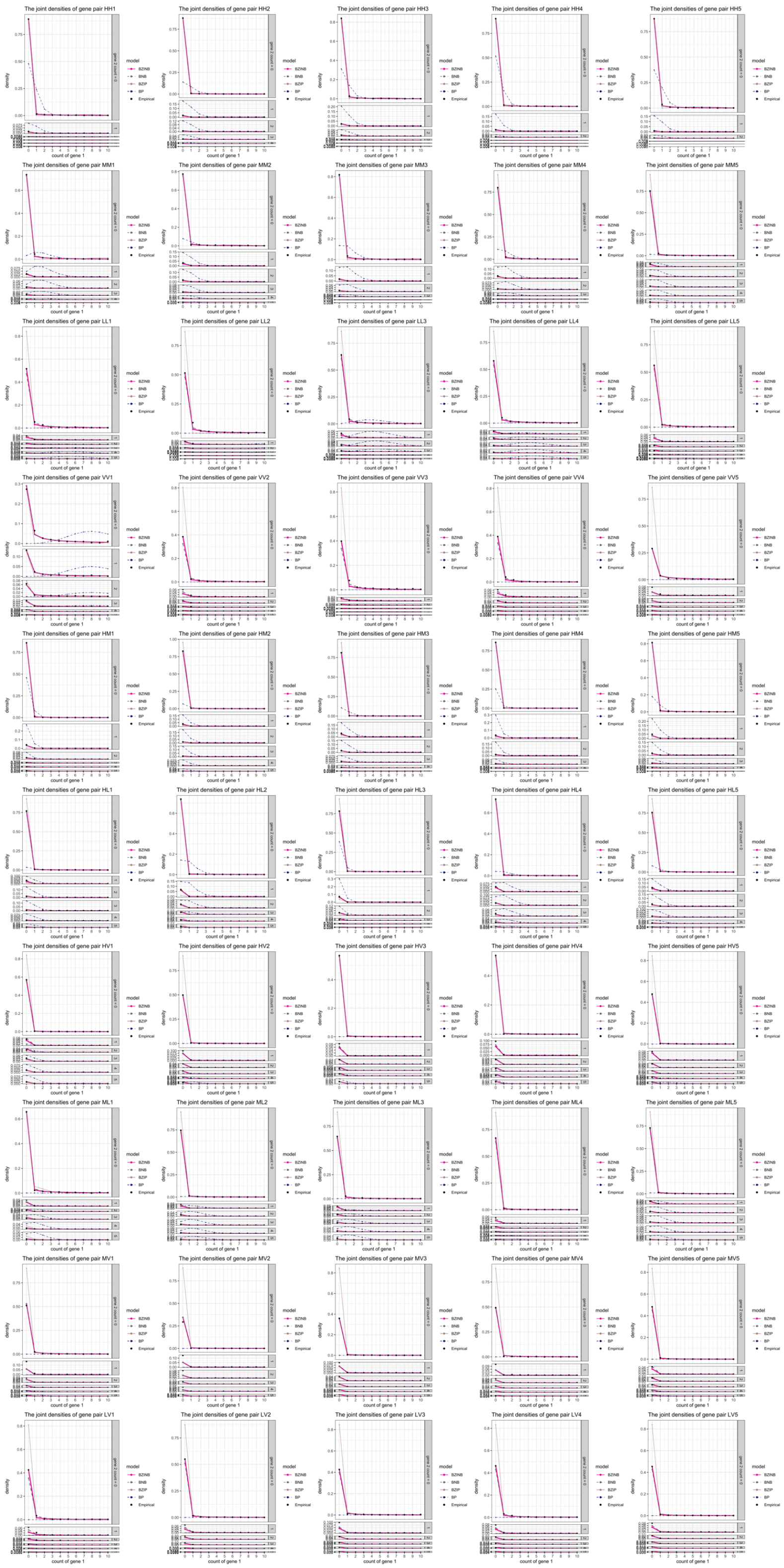


Fig. 4. The model estimates of bivariate densities (lines) and the empirical densities (dots) of all 50 gene pairs.

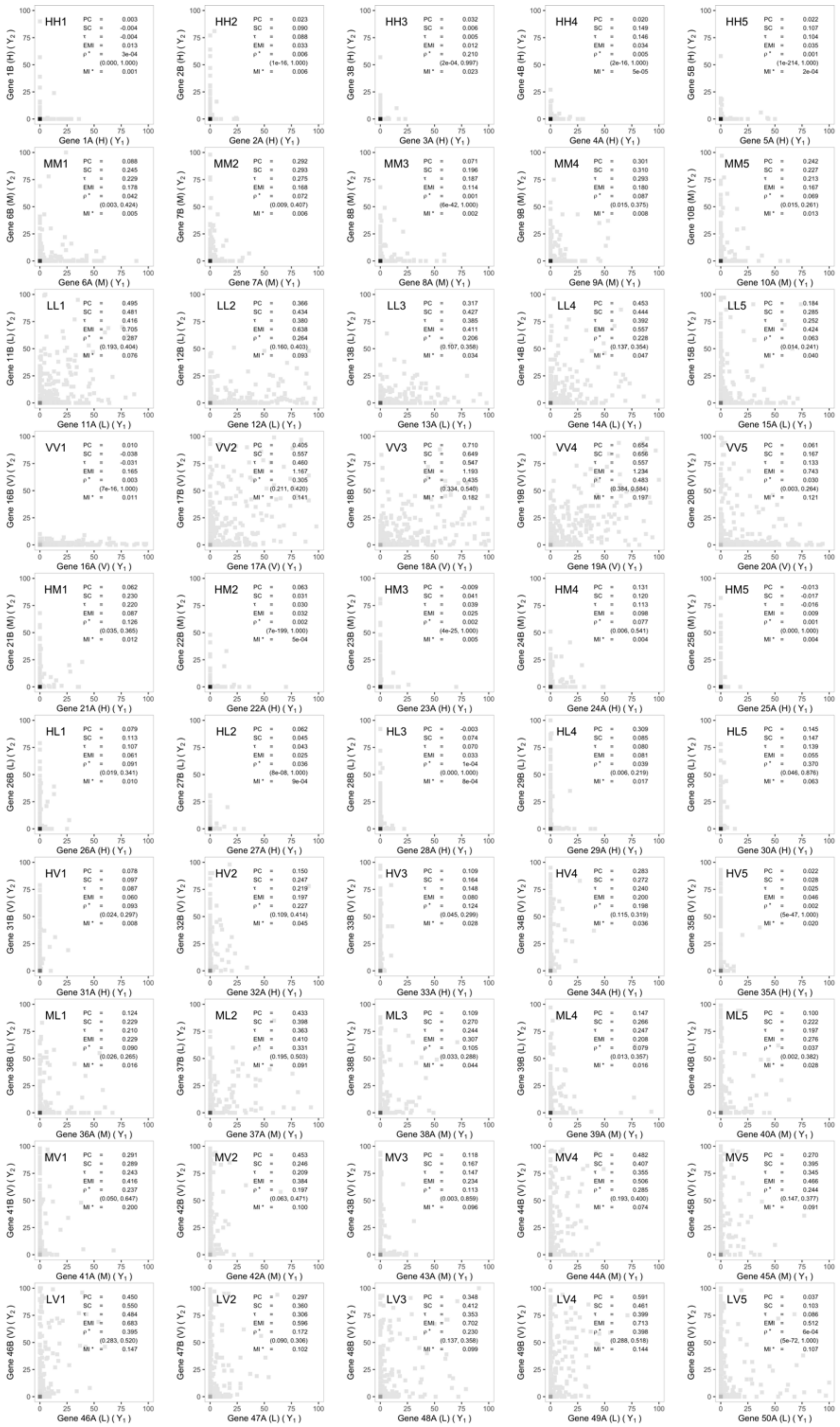

Fig. 5. The dependence measures of all 50 randomly selected pairs of mouse paneth RNA count data. Each row contains five pairs: From top to bottom, rows represent HH, MM, LL, VV, HM, HL, HV, ML, MV, and LV, in the order. For each pair, the measures and 95% confidence intervals of the $\hat{\rho}^*$ in the parantheses are presented on top of the empirical distribution plot.
